# Naturalistic Audiovisual Stimulation Reveals the Tonotopic Organization of Human Auditory Cortex

**DOI:** 10.1101/2021.07.05.447566

**Authors:** Nicholas Hedger, Tomas Knapen

## Abstract

Despite the importance of audition in spatial, semantic, and social function, there is no consensus regarding the detailed organisation of human auditory cortex. Using a novel application of a population receptive field model to a high-powered naturalistic audiovisual movie-watching dataset, we simultaneously estimate the basic spectral tuning properties and category selectivity of human auditory cortex. This revealed unprecedentedly clear tonotopic maps which showcase the modes of organization and computational motifs of the auditory cortex. Specifically, we find that regions more remote from the auditory core exhibit more compressive, non-linear response properties with finely-tuned, speech selectivity in low frequency portions of their tonotopic maps. These patterns of organisation mirror aspects of the visual cortical hierarchy, wherein tuning properties progress from a stimulus category-agnostic ‘front end’ towards more advanced regions increasingly optimised for behaviorally relevant stimulus categories.

## 1. Introduction

Auditory information processing is fundamental to human spatial, semantic, and social function. The principal dimension in which auditory sensations are organized is sound frequency. The cochlea ^1^, brainstem ^2^, thalamic ^3,4^, and cortical ^5^ centers of the auditory system all exhibit *tonotopy*: neighbouring neurons with similar spectral sensitivities give rise to a smooth topographic progression of frequency tuning within each region. The primacy of tonotopy as an organizing principle in both the auditory brain ^4,5^ and artificial neural networks of audition ^6^ is uncontested, but there is little consensus on the cortical auditory system’s tonotopic structure ^7,8^.

In analogy to the auditory brain, many visual brain regions are organized retinotopically: according to the structure of their sensory organ, the retina ^9,10^. In contrast to audition’s elusive organization, vision science has been able to leverage explicit computational models of the brain’s encoding of visual space to chart multiple distinct retinotopic maps, with the proposed number of these maps rapidly increasing in recent years. ^11–13^. The combination of retinotopic mapping with charting selectivity for salient stimulus categories ^14^ (e.g. faces, objects and environmental scenes) has delivered a relatively detailed understanding of the layered topology of stages that underlie the visual system’s processing hierarchy. In this work, our goal was to apply the computational neuroimaging toolkit of vision to the auditory system in order to reveal its topographic organization.

Originally applied to visual cortex^10^, a population receptive field (pRF) model is a parsimonious encoding model that explains population neural responses as the interaction between a stimulus and a receptive field. Previous work has leveraged this model to reveal tonotopic maps derived from pure tone stimuli^15^. The application of other computational models to explaining responses to naturalistic sound stimuli has also had great explanatory power in revealing core aspects of auditory cortical function ^16–18^. However, although such studies employed real-world stimuli, they were brief (<10s), isolated and contextless sounds that do not closely resemble everyday auditory experience. Our natural sensory experiences are much better characterised by extended and continuous multisensory presentations - much of the meaning we infer is from stimulation sequences rather than from instantaneous stimuli. In this study, we develop previous work via the novel application of a nonlinear pRF model to explain responses to more naturalistic stimuli. Moreover, we perform our modeling on a volume of data that far exceeds that underlying previous tonotopic mapping studies, offering us a unique insight into auditory cortical function.

Here, we used naturalistic movie stimuli to simultaneously chart both low-level tonotopic structure as well as higher level category-selective structure of auditory cortex. To this end, we leveraged a uniquely high-powered 7-Tesla functional imaging dataset acquired in the Human Connectome Project (HCP), in which 174 participants viewed approximately 1 hour of naturalistic audiovisual movies ^19,20^. We analyzed these data with the compressive spectral summation (CSS) model - a nonlinear pRF model that explains the hemodynamic response of cortical locations as resulting from the interaction between an auditory stimulus and the basic spectral tuning of their population receptive field (**Figure 1**, **Methods**).

**Figure 1.**
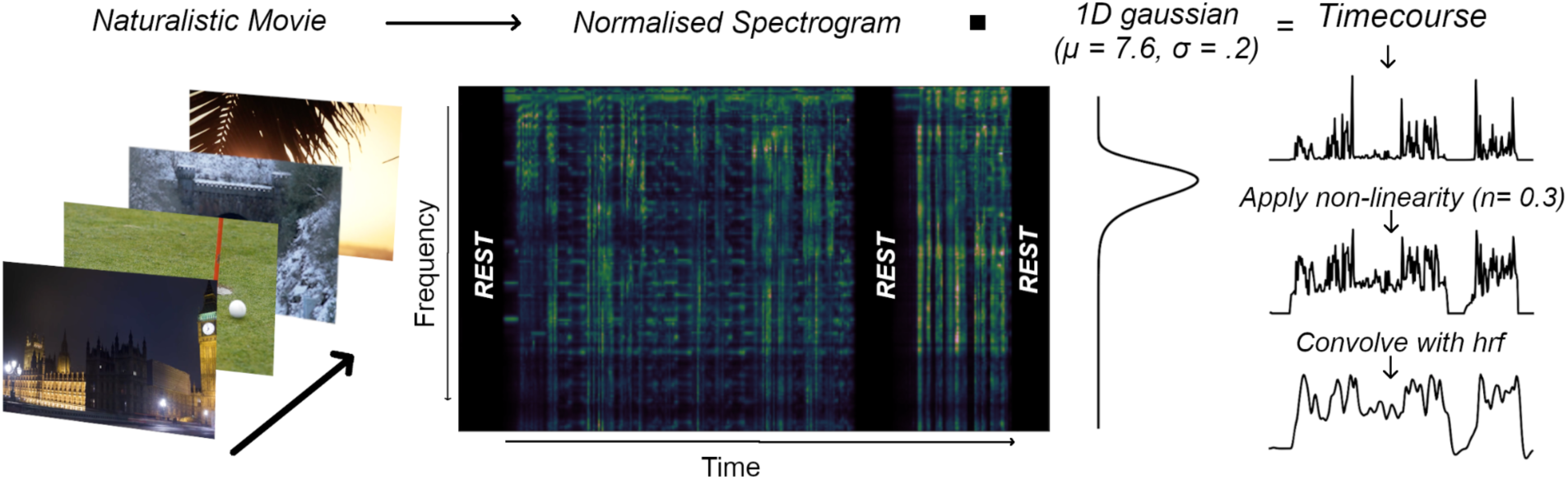
Modeling Approach. Participants were presented with an audiovisual movie. To create a design matrix for our modeling, we extracted the audio track of this movie and decomposed it into a spectrogram that represents the power spectral density of the audio signal. The spectrogram for a 5 minute segment of a movie is shown. A predicted timecourse is generated by taking the dot product of the spectrogram and a Gaussian function defined over log frequency (Hz), parameterised by its peak position (μ) and size (σ). Subsequently a static nonlinearity (n) is applied to the timecourse before convolution with a hemodynamic response function. Representative model fits, as well as the effect of varying each parameter on model predictions are depicted in **Supplementary Figure S1**.

## 2. Results

### 2.1. Global Tonotopic Organisation

To focus our analysis on regions sensitive to structured auditory information, we initially removed vertices that responded unselectively to sound (**see Methods**). This revealed a tonotopic population of vertices in a large expanse of superior temporal cortex. Preferred frequency (μ) of this population is depicted in **Figure 2A** and **2B**. The population covers Heschl’s gyrus (HG) - a landmark traditionally associated with the auditory core ^21^ and spans portions of the planum temporale (PT), superior temporal gyrus (STG) and terminates ventrally close to the superior temporal sulcus (STS).

**Figure 2.**
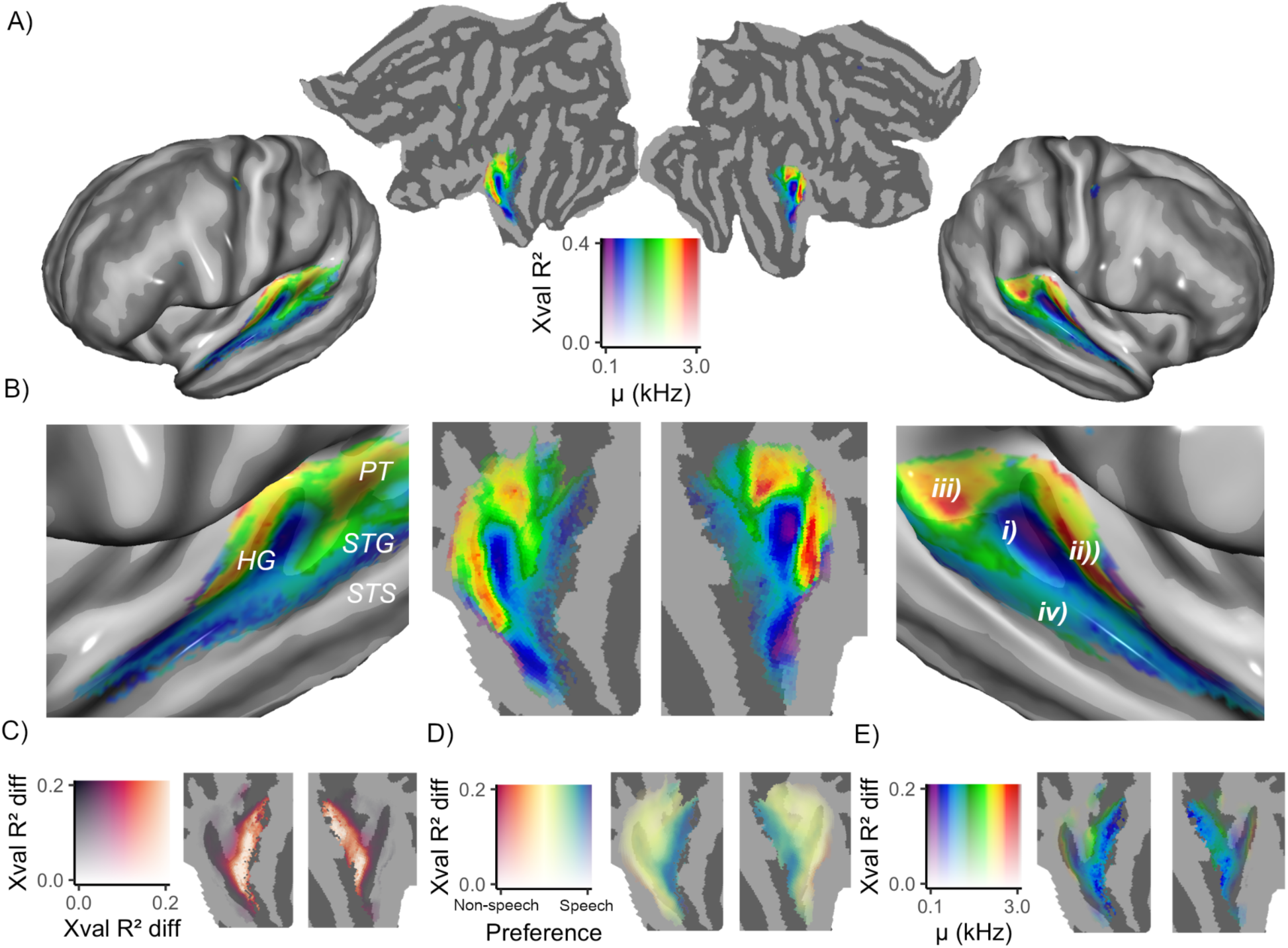
**A)** Shows preferred frequency (μ) of the tonotopic population of vertices that were designated for further analysis (see **Methods**) rendered onto flattened and semi-inflated representations of the cortical surface. The 2D colormap depicts μ across the x axis and generalisation performance (*Xval R^2^*) defined by transparency across the y axis. **B)** Depicts zoomed-in views of the data from **A**. **C)** Depicts the difference in out-of-sample performance (*Xval R^2^ diff*) between the speech-selective and CSS models. **D)** Depicts category preferences (speech v nonspeech). **E)** Depicts the μ estimated by the CSS model weighted by model improvement offered by the speech-selective model. Primarily low-frequencies near STG are visible, indicating that speech selectivity is observed at low-frequency portions of non-primary tonotopic maps

The tonotopic structure depicted in **Figure 2B** mirrors classical hallmarks of auditory organisation that have been observed in traditional psycho-acoustic paradigms ^21,22^. There is a large, ‘core’-like, low-frequency region on HG (**i**), surrounded by an antero-medial ‘strip’ (**ii**) and posterior ‘patch’ (**iii**) preferring high frequencies. These high-frequency regions adjoin approximately medially, creating the hallmark ‘V’ shaped pattern that has been observed consistently in previous investigations ^8^. As model performance drops off dorsolaterally, an additional region preferring low frequencies can be observed that roughly aligns with STG (**iv**). The organisation of these features are highly consistent across hemispheres. Although we searched for preferred frequencies up to 8 kHz, we found little selectivity for frequencies above 4 kHz. This range of frequencies revealed by our analyses are in good agreement with a complementary, high-field study based on brief, naturalistic stimuli ^16^. Moreover, in addition to the similar range of frequencies, we also note the striking similarity between the finer-level tonotopic arrangement revealed by our analyses. The range of frequencies likely reflects a combination of the fact that low frequencies tend to dominate natural sounds and the effect of across-subject averaging.

Representative model fits, as well as an assessment of the CSS model’s predictive capabilities referenced against data reliability can be found in **Supplementary Material S1 and Figure S1**. In terms of cross-validation performance, the CSS model outperformed a model with a fixed nonlinearity (n) in nearly all locations of the tonotopic population, indicating the benefit of this aspect of the modeling (**Supplementary Material S1**).To quantify the reliability of parameter estimates, we assessed the association between the μ estimates obtained from two across-subject folds (the ‘*early*’ and ‘*late*’ subject - see **Methods**). There was robust agreement between these tonotopic maps (Spearman’s *r_s_* = .87, *p* <.001) and similar levels of agreement with respect to the maps of the pRF size - as defined by full width half maximum (FWHM - *r_s_* = .86, *p* <.001) and n parameters (*r_s_* = .85, *p* <.001) . We further investigated the stability of tonotopic maps in two additional ways, performing model fitting on both individual subjects and on fold-wise and subject-wise split-halves of the dataset (**see Methods and Supplementary Material S2**). After ranking individual subject maps by out of sample variance explained, a generally consistent tonotopic arrangement was revealed in the top 5 subjects. These patterns, however, were less conspicuous in the middle 5 subjects and virtually no tonotopic arrangement was discernible in the bottom 5 subjects (**Figure S2A**). However, the split-half analyses revealed stable parameter estimates, both at across-subject and across-fold splits of the data (**Figure S2B -E**). Together, this illustrates that a large amount of data/high signal to noise ratio is required to recover the tonotopic organization we report - possibly accounting for why smaller-scale studies often yield inconsistent conclusions about tonotopic organization.

### 2.2. Global Speech Selectivity

We characterised category selectivity via a *speech-selective* model, derived from a linear combination of the CSS-predicted responses to speech and nonspeech stimuli (see **Methods**). **Figure 2C** depicts the improvement in generalisation performance for the speech-selective model over the initial CSS model. Notably, there is very little improvement around HG, but robust improvements in performance can be observed around STG. Speech selectivity (defined by difference between speech and nonspeech beta-weights) increases ventro-laterally along an axis that roughly aligns and overlaps with STG (**Figure 2D**). **Figure 2E** depicts the μ parameters estimated by the CSS model, with transparency weighted by speech-selective model improvement. Interestingly, this reveals that speech-selective regions primarily occupy the same low-frequency portions of the tonotopic map along STG that were associated with a decline in CSS performance (i.e. region **iv in 2B**).

### 2.3. Decomposition of Functional and Anatomical Parameters

The preceding modeling has quantified basic spectral tuning properties, as well as category selectivity in auditory cortex. Spectral tuning has been consistently linked to anatomical and myelo-architectural properties. For instance, the auditory core has been linked to a highly myelinated region on HG ^23,24^ and surface curvature has been linked to frequency tuning ^21^. Furthermore, delineation of auditory fields based solely on functional data remain controversial ^7,25^. It is therefore important to consider how our functional quantifications relate to anatomical and myelo-architectural properties. A succinct, hypothesis-free way of revealing the low dimensional structure of these relationships is via a principal components analysis (PCA), which can reveal the spatial organization of otherwise hidden response dimensions.

Our estimated functional parameters (the CSS parameters μ, FWHM, n, as well as speech selectivity) are depicted in **Figures 3A-D** alongside structural parameters of the HCP data (i.e. myelin density, cortical thickness, sulcal depth and curvature) in **Figures 3E-H**. We decomposed this wide set of parameters into its low-dimensional structure via weighted PCA ^26^. The first 3 components, which together accounted for 83% of the variance explained and showed a high degree of consistency across hemispheres are illustrated in **Figures 3I-M**. The feature loadings for each component are summarised in **Figure 3I** above the components themselves in **Figures 3J-M**.

**Figure 3.**
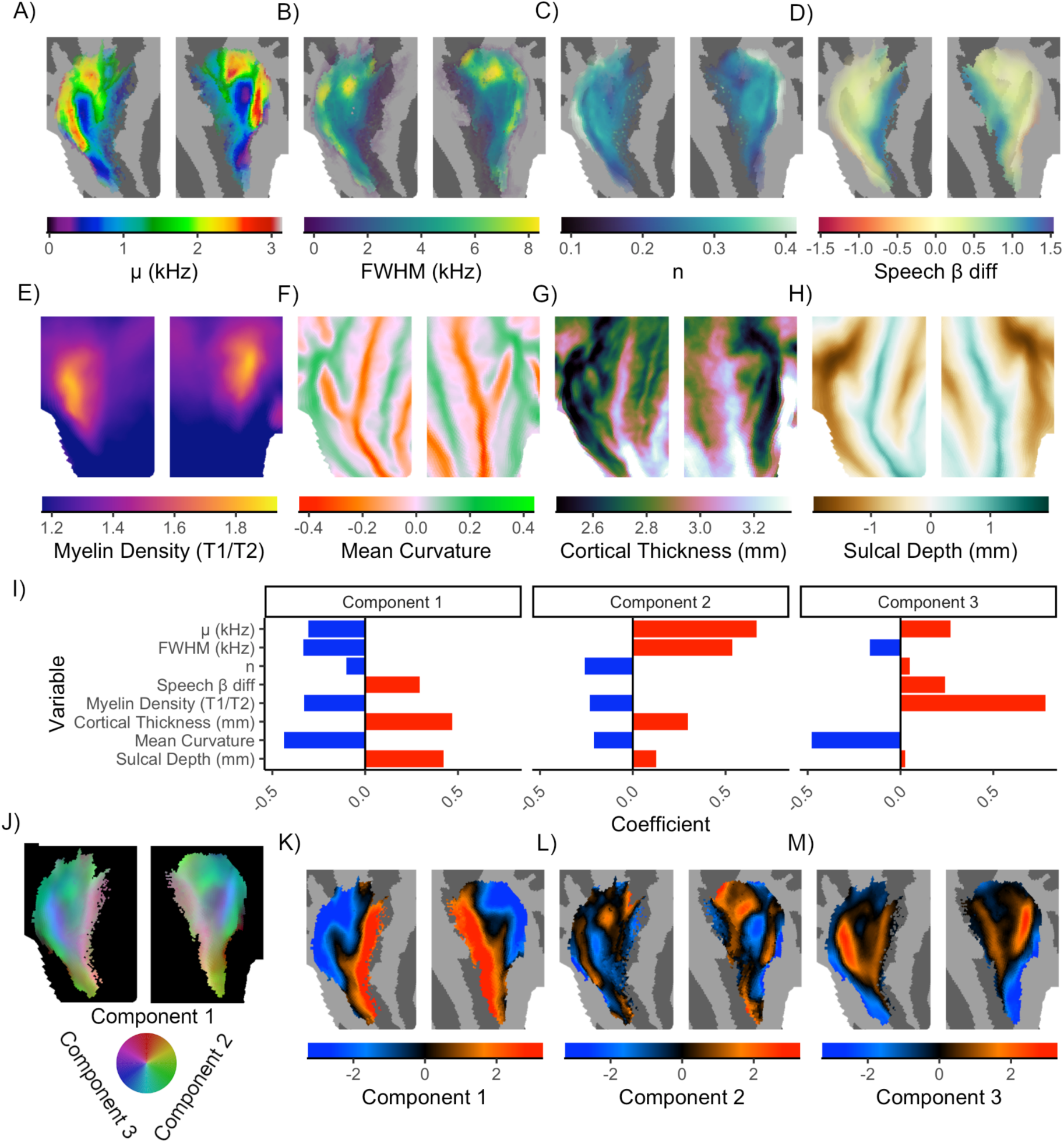
Panels **A-D** depict functional parameters of the data derived from the CSS and speech selective models. Panels **E-H** depict averaged statistics for myelo-architectural and structural parameters. **I)** Depicts the feature loadings onto each of the 3 principal components. **J)** Shows the 3 components overlaid in RGB color space, as depicted by the color-wheel. Panels **K-M** depict the individual components.

The PCA components reveal separable modes of organisation, ranging from spatially broad to intricate, that differ substantially in their relations to structural and functional parameters. Component 1, for instance, reflects a broad pattern of spatial organisation mostly tied to the structural features, increasing in a roughly postero-medial to antero-lateral direction around the medial tip of HG to STG (**Figure 3K**). Functionally, it also reflects a tendency for μ, FWHM and n to decrease and speech selectivity to increase towards STG. We note that the spatial profile of this component is broadly consistent with that revealed by a complementary hypothesis-free decomposition of auditory cortex, wherein acoustically driven responses were observed towards HG, whereas voxels whose response poorly explained by frequency statistics and better explained by speech-selectivity were observed towards STG (see component 5 of ^27^). Component 2 (**Figure 3L**) which has a clear resemblance to the preferred frequency data in **Figure 2B**, exhibits a more intricate structure that is most strongly tied to the parameters estimated by the CSS model. In particular, it depicts a region on HG characterised by low μ and FWHM surrounded laterally and medially by regions preferring higher frequencies with more compressive responses. This component illustrates well the previously observed tendency for higher frequencies to be observed at increased sulcal depth ^21^ - as well as for high frequency regions to be associated with larger bandwidths ^28^. Moreover, it also depicts increased non-linearity (lower values of n) in regions surrounding HG, consistent with the greater nonlinearity observed in non-primary sensory regions of the visual cortex ^29^. Component 3 (**Figure 3M**) depicts a prominent, heavily myelinated region on HG flanked medially by high FWHM and curvature. Since increased myelin density has been linked to the location of primary sensory cortices, including human A1 ^23^, and myelination here is associated with lower curvature, we interpret the spatial profile of this component as being consistent with presence of a primary ‘core’ region that runs along HG. These patterns of organisation revealed by the PCA were also highly consistent in the ‘early’ and ‘late’ subjects (**Supplementary Figure S3**).

### 2.4. ROI-based Quantification of Functional Properties

To provide spatial precision to our quantifications, we next defined a set of regions of interest (ROIs) based on the presence of tonotopic reversals ^22,30^, as well as variations in tuning width ^28,31^ (here defined as FWHM), speech selectivity ^27,32^ and myelin density ^33^ (see **Methods** for details and alternative ROI definitions). The resulting scheme, comprising of 3 ‘core’ ROIs (*A1, R, RT*) and 4 ‘belt’ ROIs (*MBelt, LBelt, P1 and P2*) is depicted in **Figure 4A-B**. To provide a simplified visual representation of the tonotopic gradients present in the data, we fit a linear model to the μ data in each ROI, with the cartesian coordinates of the flattened cortical surface as regressors. These predictions, depicted in **Figures 4A-4B** capture the tonotopic structure of the data very well (*R*^2^ = .71, *B =.99, p* < .001).

**Figure 4.**
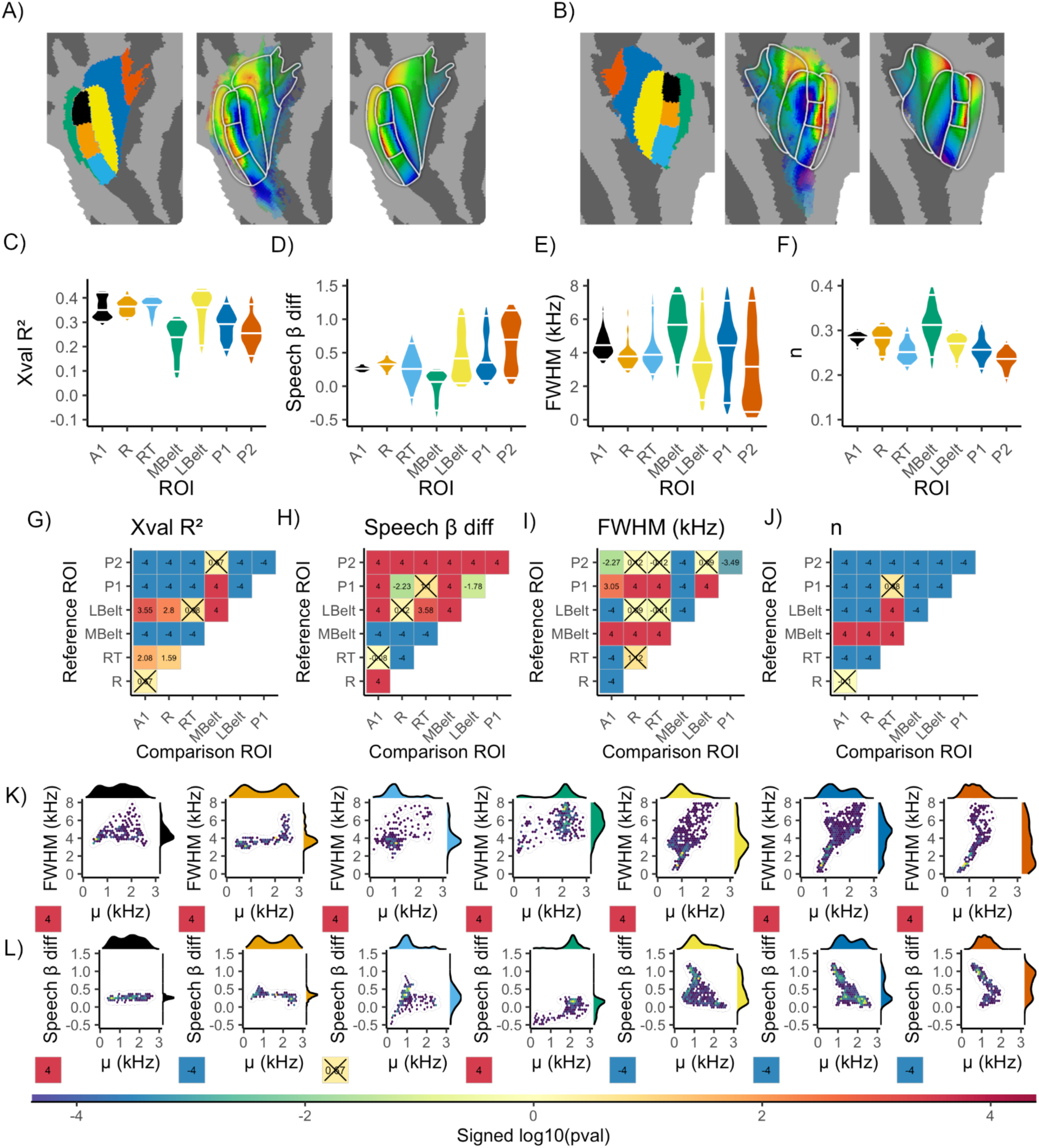
Panel **A** (left to right) shows the ROI definitions, with ROI color corresponding to the colours of the violin /marginal density plots in the remaining figures (left), the μ data (middle) and the linear fit to the μ data per ROI (right) for the left hemisphere. Panel **B** shows the same for the right hemisphere. Panels **C-F** show the per ROI distributions for out-of-sample variance explained by the CSS model (**C**), speech selectivity (**D**), FWHM (**E**) and n (**F**). Violins are normalised to have equal maximum width. The 5th, 50th and 95th percentiles of the distributions are demarcated by white horizontal lines. Panels **G-J** depict the between-ROI pairwise differences for each of the corresponding parameters in panels **C-F**. The values in each cell are signed log10 transformed *p* values. Larger values for a parameter in a reference ROI (y axis) compared to a comparison ROI (x axis) are assigned positive values (p values are -log10 transformed). Smaller values in the reference ROI are assigned negative values (*p* values are log10 transformed). *p* values that do not reach the false discovery rate corrected alpha level of .05 are marked by a cross in the corresponding cell. Panel **K** shows hexbin plots for the relationship between μ and FWHM in each ROI. The cells at the bottom left of each plot depict the same signed log10 transformed *p* values (now with positive values signifying positive relationships). Panel **L** shows hexbin plots for the relationship between μ and speech selectivity. The corresponding colorbar for all *p* values can be found at the bottom of the figure.

With ROIs established, we next examined their functional properties. The per-ROI distributions of functional parameters are depicted in **Figures 4C-F**. Between-ROI pairwise differences in these parameters were assessed via weighted-bootstrap procedures (**see Methods**). Since this results in 21 pairwise tests per parameter (with the majority revealing detectable differences), we restrict this section to a brief summary of the salient patterns in the data. The *p* values associated with each pairwise difference are depicted in **Figures 4G-J**.

In terms of variance explained by the CSS model (**Figure 4C**) the poorest model performances are observed in ROIs occupying the peripheral extremes of the tonotopic population (*MBelt, P1 and P2*), likely reflecting the principle that responses more remote from primary sensory regions will be less yoked to the low-level sensory properties of the stimulus. The ROIs vary substantially in speech-preference (**Figure 4D**) with the highest speech preferences observed laterally to the core (*LBelt, P1, P2*). The FWHM parameter was associated with the least robust differences between ROIs: there was a relatively small range of FWHM values in core regions, but substantial ranges of low and high values in non-core ROIs (**Figure 4E**). This pattern entailed that all core ROIs had detectably lower FWHM relative to *MBelt*, but were not detectably different from, or were larger than ROIs lateral to the core. The most robust differences between ROIs were detected for the exponent parameter n (**Figure 4F**). Responses became more compressive with increased distance laterally from the core across *LBelt, P1 and P2*. More medially, *A1, R* and *MBelt* had higher n values than all other regions.

Previous studies of auditory cortex have indicated that preferred frequency is positively associated with tuning bandwidth ^28,34^, indicative of a Weber’s-law type effect that mirrors the relation between pRF size and eccentricity observed in visual cortical field maps ^9^. We observed that the scaling of FWHM with μ varied systematically across ROIs, with increasingly positive relationships observed in regions more lateral to the core (**Figure 4K**).

Finally, since our initial inspection of the speech-selective model outcomes implied spatial anisotropies in the relationship between preferred frequency and speech selectivity (**Figure 2F**), we formally characterised this relationship within each ROI (**Figure 4L**). In core regions, speech selectivity is low and relatively independent of μ, but this relationship becomes strongly negative in regions lateral to the core. This pattern of results validates our observation that speech-selective regions tend to occupy low frequency portions of the tonotopic map, specifically those residing outside the core.

## 3. Discussion

Drawing inspiration from computational models of vision, our analysis elucidates the computational motifs and modes of organisation that underlie naturalistic auditory processing in human cortex. Previous studies have generated a somewhat unclear and inconsistent picture of the finer level tonotopic organisation in auditory cortex. In part, this can be attributed to diversity in methodological and analytic variables such as scanner field strength ^16^, complexity of stimuli ^28^, modeling approach^35^, spatial smoothing^36^ and criteria for designating voxels for analysis^37^. Beyond these, a large contributor is the simple limitation in the quantity of data analysed, both in terms of the number of subjects and the data per subject. To our knowledge, our modeling exploits a volume of data that far exceeds previous tonotopic mapping studies, implying an unprecedented level of statistical power. Here, we observed tonotopic reversals along HG, consistent with the tripartite A1/R/RT field organization proposed in other primates ^38^. Furthermore, we observe structured tonotopic organization both lateral and medial to HG - allowing us to define multiple cortical gradient maps beyond traditionally defined ‘core’ regions. As such, our data provides a working model of human auditory organization beyond the core regions A1 and R.

Speech selectivity was primarily observed in close proximity to STG, consistent with the long-established cortical site attributed to speech processing ^39^. Our data extend these findings by quantifying the basic spectral tuning properties of both these and more primary regions, clarifying the computational motifs that characterise the transition from acoustically-driven to speech-selective auditory cortex. Our quantifications are in excellent agreement with a recent hierarchical model of human speech processing that proposes a medial to lateral sequence of ‘spectral’ to ‘articulatory’ to ‘semantic’ representations that progress from HG to STG to STS ^40^. Here, we reveal a similar trajectory reflected in the tuning properties of a sequence of medially to laterally oriented tonotopic maps. In A1, we observed *n* values very close to .3, a biologically interpretable value for the first auditory cortical site, since it is consistent with the degree of amplitude compression performed by the input sensory organ - the cochlea ^1^. From A1, there was a medial to lateral gradient of increased compressivity, consistent with the proposed ‘cascade’ architecture of the visual hierarchy - wherein later regions add nonlinearities to the outputs from earlier regions ^41,42^. In vision, such nonlinearities have functional benefits - allowing the responses of face or object selective neurons to tolerate stimuli that appear in different sizes or locations of their receptive field ^43^. Considered with previous observations, our data indicate that similar mechanisms may support flexible tolerance of salient auditory ‘objects’ such as vocalisations. For instance, Marmoset data indicate that spectrally nonlinear responses in ‘harmonic template neurons’ may be optimised for encoding vocalisations ^44^. However these analyses were restricted to the auditory core, preventing conclusions about the trajectory of spectral nonlinearity across auditory fields. Moreover, the translatability of such a proposal based on these data is equivocal; complex, pitch-sensitivity is instead primarily evidenced outside core regions in humans ^45,46^. Here, we explicitly modeled spectral nonlinearity and speech selectivity throughout auditory cortex, revealing a shared medial to lateral trajectory that may reflect the hierarchical transition from low-level spectral representations to flexible phonemic representations that are relatively invariant to acoustic differences. Such inferences about more complex aspects of receptive fields highlight the utility of modeling compression at the cortical level, not just the input stage.

In addition to the spectral nonlinearity quantified here, there is evidence for temporally nonlinear responses in auditory cortex - BOLD responses to long-duration stimuli are less than predicted by the linear prediction from briefer duration stimuli ^47^. Studies of visual cortex have indicated increased temporal subadditivity in higher-level regions, mimicking the profile of spatial compressivity ^48^. It is possible that a similar shared trajectory of spectral and temporal compressivity also underlies responses in auditory cortex. As such, an important goal for future work is to develop a space-time model that interrogates both spectral and temporal nonlinearities.

We observed increasingly finely-tuned, speech-selective receptive fields in low frequency portions of more lateral tonotopic maps. This implies the cortical magnification of spectral features diagnostic of speech and natural soundscapes. By extension, it may reflect information-theoretic efficient coding principles, as illustrated by the observation that learned kernels optimised for encoding speech predict the positive frequency-bandwidth association observed in cochlear filters ^49^.

The close relationship between category and spectral selectivity would seem to point more to the proposed non-hierarchical structure of auditory cortex ^50^, but similar phenomena also characterise high-level sites of visual cortical hierarchy. The fusiform face area (FFA) has a combined preference for faces and central visual field locations, whereas the parahippocampal place area (PPA) prefers environmental scenes and peripheral visual field locations ^51^ - links between topography and category-selectivity that reflect their computational goals. It is therefore revealing that speech selectivity was relatively independent of preferred frequency in core regions - whereas this relation was strongly negative in regions lateral to the core. This anisotropy in tuning properties agrees with the auditory processing hierarchy implied by recent neural network models, that progresses from a ‘front end’ that performs a relatively stimulus category-agnostic analysis of arbitrary sounds towards more advanced regions that are increasingly optimised for salient sound categories, such as speech ^6^. It is important to note, however, that natural sounds contain correlations between low and higher-order features - such as the covariation of speech sounds and low frequency energy modulations. As such, the low frequency tuning estimated in speech-selective regions could reflect genuine spectral tuning, or a simple epiphenomenon of higher responses to speech sounds - creating a circular impasse. One powerful approach to address this issue is to compare responses to natural and ‘model matched’ stimuli that decorrelate low-level and high-level natural sound statistics. Such studies have revealed large divergences in the responses of non-primary regions that were hidden by feature correlations in natural stimuli ^18^. In the present case, however, we note that low frequency tuning along STG replicates findings revealed by simple pure-tone stimuli ^15,28,36^, speaking against the idea that it reflects a simple product of higher responses to speech sounds.

It is essential to reflect on potential caveats of leveraging naturalistic stimuli for mapping unimodal responses. First, an audiovisual stimulus could be considered suboptimal for tonotopic mapping due to interaereal inhibitory responses between visual and auditory cortices ^52,53^ - heteromodal sensory data can compete for access to attentional and memory resources ^54^. However, in more naturalistic contexts, cross-modal stimulation bestows many benefits - particularly in humans. For example, synchronous audiovisual presentations have been found to increase both the reliability and precision of responses in auditory cortex ^55,56^. The ability to exploit the visual source of auditory information (for instance, a speaker’s mouth) provides rhythm and amplitude information that assists a listener in directing attention towards the relevant auditory envelope, thereby boosting sensory responses ^57,58^ . Such phenomena concord with the notion that merging the senses results in a more robust cross-modal percept that augments auditory scene analysis ^59^. Moreover, despite crossmodal auditory influences on visual cortex, robust retinotopic maps have been revealed from the same HCP movie-watching data that are highly consistent with those derived from unimodal visual stimulus presentations ^12^. These considerations, together with the consistency of the principles we reveal with those derived from unimodal auditory stimulation, highlight the utility of naturalistic movie stimuli for revealing principles of auditory organization.

Second, to enhance signal to noise ratio, our central analyses were conducted on across-subject averaged data. As such, they do not capture the idiosyncrasies of individual subjects - which are particularly prominent in non-primary areas. Despite this limitation, increasing the SNR in this way yields tangible benefits. This is illustrated by the outcomes of individual subject analyses (**Supplementary Material S2**), which indicate that a large amount of data is required to recover parameter estimates with good generalization performance. This notion is reinforced by the recent discovery of topographic organization in both the cerebellum ^60^ and hippocampus ^12^, which can be attributed to the unprecedented SNR offered by extensive across-subject averaging afforded by large, carefully aligned datasets such as the HCP dataset. Using traditional anatomical co-alignment methods, across subject averaging would risk regression to the mean via neighbouring tonotopic regions merging together, due to the individual variability of cortical field map sizes and positions ^7^. As such, the vivid tonotopic arrangement we observe speaks both to the robustness of tonotopy as an organising principle and also to the robustness of the HCP cross-subject alignment. Critically, the HCP data were aligned by leveraging ‘areal features’ (myelin and resting state connectivity maps) explicitly optimised to align primary sensory cortices ^61,62^, from which our analyses will have benefited greatly relative to traditional alignment based on cortical folding patterns. We must recognise, however, that alignment of regions positioned remotely from primary areas via this method is likely less optimal. This, combined with our conservative criteria for selecting vertices for analysis (see **Methods** and **Supplementary Material S6**), possibly impeded detection of subtle tonotopic arrangements in some non-primary regions. For instance, inconsistent with one recent phase-encoding study ^36^ the dorsal fields of the STS effectively demarcated the ventral boundary of tonotopic responses (**Supplementary Figure S5)**. The speech-selective tonotopic portions of STG may represent initial stages of speech processing wherein the spectral properties of speech sounds begin transformation into more abstract representations ^28^. The neighbouring STS, which exhibits complex multi-modal responses with little visuo-spatial selectivity ^63^ or spectral selectivity, may represent a site wherein socially relevant features are coded from abstracted forms of audiovisual signals ^64^.

Our study also reveals some interesting implications for estimating pRF size. First, psychophysics experiments allow stimulation to be optimized for estimating pRF size across a range of frequencies, whereas naturalistic stimulation limits this range. The frequency profile of naturalistic soundscapes may therefore lead to an underestimated pRF size at higher frequencies. Secondly, pRF size is conceptually difficult to define when the underlying system exhibits the compressive behavior observed here and linear models typically overestimate pRF size in such cases ^29^. We compensate for this by normalising by the exponent and estimating pRF size by simulating responses to infinitesimally punctate stimulation (see **Methods**) - but this cannot circumvent the inherent interaction between stimulation extent and pRF size implied by compressivity. Common to all fMRI studies is the limitation that voxels containing neurons with diverse preferred frequencies (e.g. along HG) would likely be estimated to have larger pRF sizes than regions with homogenous preferred frequencies (e.g. along STG). Lastly, whilst we capture more complex response profiles than a linear model, estimates of pRF size only reflect the width of a main spectral peak. It has been demonstrated that portions of nonprimary auditory cortex exhibit sensitivity to multiple frequency bands at an octave distance from one another, harmonically related intervals, or with no clear relation ^31,44,45^. As such, our measure of pRF size approximates the width of a simple population response function - likely reflecting something fundamentally different from ‘bandwidth’ as estimated from single cell recordings. Together, these difficulties may account for the inconsistent frequency-bandwidth relations observed in human fMRI studies, with some indicating principled relationships ^28^ and others revealing no clear relationships ^15,46^.

We must also consider disjunctures that heavily constrain the analogies we draw between vision and audition. Firstly, retinotopic mapping models a mobile sensory array that samples locations of the visual field, whereas tonotopic mapping models the spectral signature of auditory signals largely incommensurate with their spatial location. This difference likely explains why pRF sizes are reliably larger in more advanced regions of visual cortex ^10,29^, whereas a range of narrowly and broadly tuned pRFs can be found outside the auditory core ^15,28^. Effective processing of objects, faces and scenes are facilitated by location invariance and pooling signals from large extents of the initial sensory space ^29^. Conversely, pooling large extents of frequency space could impede high level tasks such as speech processing, since speech sounds tend to occupy restricted and predictable portions of the sensory space (primarily low frequencies).

Another divergence is that primary auditory cortex would also occupy a later position in a processing hierarchy than its visual counterpart, since there is a more extensive sequence of auditory than visual nuclei in subcortex ^1,2^ with comparatively later layers of neural networks best explaining responses in primary auditory cortex ^6^. Information in ‘primary’ auditory regions thus represents a relatively processed format of the initial sensory data. Accordingly, additional explanatory power could be derived from modeling other principled dimensions of auditory data, such as periodicity ^65^, selectivity for stimulus categories such as music ^27^ and spatial cognition or explicit recognition task contrasts ^66,67^.

To summarise, via a novel application of a nonlinear population receptive field model to naturalistic stimuli, we elucidate the detailed topography of human auditory cortex. This clarified the major sensory reference frame that underpins higher level auditory cognition, and revealed computational motifs that mirror classical hallmarks of the visual system. Such similarity may reflect common organisational principles shared across multiple sensory systems, aiding the discovery of novel topographic organisational structures in the representation of other senses.

## Methods

### Participants and Stimuli

Data were taken from the 174 participants of the HCP movie-watching dataset ^19,20^. The sample consisted of 104 females and 70 males (*M* age 29.3 years, *SD* = 3.3) born in Missouri, USA. 88.5% of the sample identified as ‘White’ (4.0% ‘Asian’, ‘Hawaiian or Other Pacific Island’, 6.3% ‘Black or African American’ 1.1% unreported). The English language comprehension ability of the sample (as assessed by age-adjusted NIH Picture Vocabulary Test^68^ scores) was above the national average of 100 (*M* = 110, *SD* = 15).

Participants were scanned while watching short (ranging from 1 to 4.3 minutes in length) independent and Hollywood film clips that were concatenated into movies of 11.9 - 13.7 minutes total length. Before each clip, and after the final clip was displayed, there were 20 second ‘rest’ periods wherein there was no auditory stimulation and only the word ‘*REST*’ presented on the screen. There were 4 separate functional runs, wherein observers viewed 4 separate movies. All 4 movies contained an identical 83 second ‘validation’ sequence at the end of the movie. Audio was scaled to ensure that no video clips were too loud or quiet across sessions and was delivered by Sensimetric earbuds that provide high-quality acoustic stimulus delivery while attenuating scanner noise. Movie audio format was set to AAC-LC, 192 kbps, stereo, with a sampling rate of 44.1 kHz, entailing a frequency range of <= 22 kHz. Spectrograms for each of the movies are available in **Supplementary Material S7**. Full details of the procedure and experimental setup are reported in the HCP S12000 release reference manual.

### Data Format and Preparation

Ultra-high field fMRI (7T) data from the subjects were used, sampled at 1.6 mm isotropic resolution and a rate of 1 Hz ^19^. For all analyses, the Fix independent component analysis-denoised time-course data, sampled to the 59,000 vertex-per-hemisphere areal feature-based cross-subject alignment method (MSMAll- ^62^) surface format was used. These data are freely available from the HCP project website. The MSMAII method is optimised for aligning primary sensory cortices based on variations in myelin density and resting state connectivity maps ^61,62^ . Because of the unreliable relation between cortical folding patterns and functional boundaries, MSM method takes into account underlying cortical microarchitecture, such as myelin, which is known to match sensory brain function better than cortical folding patterns alone ^69^. Previous research has demonstrated that such an approach improves the cross-subject alignment of independent task fMRI datasets while at the same time decreasing the alignment of cortical folding patterns that do not correlate with cortical areal locations ^62^.

For the purposes of cross-validation, we ensured that functional runs sampled only unique stimulus content. We did this by removing the final 103 seconds of each movie and corresponding functional data that corresponded to the identical ‘validation’ sequence and the final rest period. For each run, BOLD time series data were then converted to percent signal change. We next applied a high-pass filter to the data. We implemented this via a Savitzky Golay filter (3rd order, 210 seconds in length), which is a robust, flexible filter that allowed us to tailor our parameters to reduce the influence of low frequency components of the signal unrelated to the content of the experimental stimulation (e.g. drift, generic changes in basal metabolism). The effect of this temporal filtering on data and model predictions is illustrated in **Figure S1E.**

We decided to create two time-course averages that reflect a pre-existing split in the HCP dataset. Specifically, each participant viewed one of two slightly different versions of the movies. As such, performing analyses on the full set of subjects required us to respect any differences in the videos and resulting design matrices. Accordingly, we created two across-participant time course averages. The *‘early’* subject was averaged across the participants that viewed the versions of the videos before August 21st, 2014 (*N* = 42). The *‘late’* subject was averaged across the participants that viewed the movie after this date (*N* = 132). In addition to respecting the minor differences in the videos that were presented, dividing the data in this way also allowed us to characterise the split reliability of parameter estimates and also assess the impact of statistical power, which differs between these two across-participant folds.

### Design Matrix

To create a design matrix for our pRF modeling, the original .mp4 file for the movies were converted to a .wav audio file and then submitted to a Fourier transform via scipy’s *‘signal.spectrogram’* function, to generate spectrograms that represent the power spectral density of the audio signal. The resulting spectrograms were then mean-downsampled into the TR resolution and normalised via division by the corresponding per-movie, per-frequency standard deviations.

### Compressive Spectral Summation pRF Model

A pRF model is a parsimonious encoding model that characterises the relationship between a stimulus and the response from a tuned population of neurons ^10^. We modeled the pRF as a 1D Gaussian function defined over log auditory frequency (Hz). The model generates a predicted response by computing a weighted sum of the pRF and the spectrogram and then applying a static power-law nonlinearity. This can be expressed formally as:

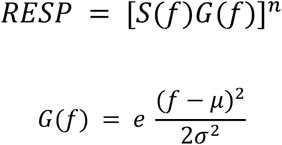

where RESP is the predicted response, *f* is the auditory frequency, S is the spectrogram, G is a Gaussian, n is an exponent parameter and μ and σ are parameters that control the mean and standard deviation of the Gaussian. The resulting timecourses (RESP) generated by the model were convolved with a double-gamma hemodynamic response function. Finally, to be conservative, and respect the same temporal filtering that was applied to the underlying data, our model predictions were also filtered with an identical Savitzky-Golay filter to that applied to the functional data. We incorporated a power-law nonlinearity in our modeling to capture a wide range of response properties with a single parameter that is straightforward to interpret. Lower values of n indicate greater compressive nonlinearity and imply that smaller amounts of overlap between the prf and the stimulus produce larger responses. In vision, these more compressive response properties are characteristic of later regions of the visual hierarchy ^43^. In application to spatial vision, this model has been referred to as the compressive spatial summation (CSS) model ^29^. Here, in its application to audition and the spectral domain, we refer to it as a compressive *spectral* summation model. In such a model, it is important to account for the fact that the effective size of a pRF is influenced not only by σ, but also by n, since nonlinearity implies that pRFs with a small σ may still respond strongly to frequencies remote from μ if n is highly compressive. A pRF’s response profile to ‘point’ stimuli placed across frequency space is therefore a Gaussian that is scaled by n. The standard deviation of this Gaussian profile can therefore be calculated by correcting for this scaling via normalisation by n.

We therefore performed the following adjustment to σ:

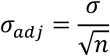

Importantly, this definition of pRF size is derived from input-output characteristics - meaning that it can be applied to any model of an auditory receptive field ^29^. To provide a final measure of pRF size, we calculated the full width half maximum of these adjusted pRF functions in kHz. Alternative definitions of pRF size expressed in octaves are presented in **Supplementary Material S8**.

### Model Fitting and Vertex Selection

#### CSS model

Model fits for each vertex in the brain were obtained by finding parameter combinations that maximized the correlation between the predicted and observed fMRI time-courses. This parameter search consisted of a coarse-to-fine strategy. We first performed an initial coarse grid search of μ and σ values, based on the initial parameter space sampled in a previous pRF study of auditory cortex ^15^ . μ was bounded between .088 kHz and 8 kHz and σ had an upper bound of 4. The best fitting parameters from this set were then used as starting parameters for a nonlinear search algorithm (scipy’s *minimize* function, using the L-BFGS-B method) which uses nonlinear minimization to find the pRF model parameters μ and σ that maximize the correlation between the pRF-predicted time-series and the observed fMRI time-course. These parameters for μ and σ were then submitted as starting parameters to a final iterative search, wherein the full set of model parameters - μ, σ and additionally the exponent parameter (n) were estimated.

To assess the generalisation performance of the pRF models, we used a leave-one-run-out cross-validation strategy. Models were fit to four separate folds of data. In each fold, the training data were concatenated across 3 movies and the fitted parameters were used to generate predictions for the left out movie. The out-of-sample performance of these predictions was then assessed by the coefficient of determination (*R*^2^). All pRF model fitting procedures were carried out using procedures from a dedicated python package: *prfpy.* To summarise the fold-wise parameter estimates, we calculated across-fold averages of the parameters in a manner that was weighted by the corresponding out-of-sample *R^2^*. This procedure was carried out for both the ‘early’ and ‘late’ subject before the parameters obtained from each were again weighted-averaged to provide final parameters.

To focus our analysis and select vertices for further analyses, we used a conservative, null-model based criterion to exclude non-tonotopic vertices (**Supplementary Material S6 and Supplementary Figure S9**). These procedures resulted in a ‘tonotopic population’ consisting of 3760 from the total 118584 cortex vertices being included in further analyses. We did not perform any spatial-smoothing of data or model parameters at any point in analysis or presentation of our data.

#### Speech-selective model

In addition to purely tonotopic responses captured by the CSS model, it is important to characterise the combination of both tonotopic and speech-selective responses. To this end, we additionally fit a simple *speech-selective* model that involved a linear combination of the separate CSS model-predicted responses to speech and nonspeech stimuli. HCP movie files were privately uploaded to YouTube. We employed YouTube’s auto-captioning algorithm to provide an initial estimate of the onset and offset times of speech sounds. These initial estimates were then manually fine-tuned by NH. Corrections mostly involved adding brief speech sounds that occured in the presence of ambient noise, or adjusting offset times, which were occasionally estimated to occur beyond the actual duration of the speech. With onset and offset times refined, we created two new design matrices. The *speech* design matrix was created by coding parts of the spectrogram overlapping with nonspeech sounds as 0. The *nonspeech* design matrix was the inverse - with all instances of speech coded as 0. We then used the parameters estimated by the CSS model to generate predictions based on these new *speech* and *non-speech* design matrices. These predictions were then entered as regressors for the design matrix of the speech-selective model, wherein we estimated the linear combination of the speech and non-speech regressors that minimised the error between the predicted and observed timecourses for every cortical location separately. Hence, this model estimated independent beta weights for speech and non-speech sounds whilst assuming μ σ and n are fixed for periods of speech and nonspeech. Generalisation performance was evaluated with the same cross-validation strategy as reported for the CSS model.

### Individual Subject Fitting

To focus our analysis of individual subjects, we restricted our model fitting to the tonotopic population of 3760 vertices derived from the aggregated data. Otherwise, identical model fitting procedures were applied as described in **Model Fitting and Vertex Selection**. To depict the stability of the tonotopic maps in individual HCP participants, we ranked individuals based on the median out of sample variance explained across this population and display data from the top, middle and bottom 5 ranked participants **(see Supplementary Figure S2A)**.

### ROI Definitions

Approaches to delineating distinct regions of auditory cortex are heterogenous, ranging from a simple, binary distinction between ‘core’ and ‘non-core’ regions ^15,27^, to extrapolating detailed non-human primate atlases onto human data ^30,37^, or defining multiple regions as remote as the STS based solely on tonotopic reversals ^36^ . This diversity may reflect the preference for ‘lumping’ versus ‘splitting’ of distinct cortical areas ^9^. To respect this range of perspectives, we used 3 strategies to define regions of interest (ROIs) within which to provide spatial precision to our quantifications. A detailed account of each of these strategies is provided in **Supplementary Material S4 and Supplementary Figures S4-S5**. A separate controversy concerns the orientation of the auditory core with respect to HG, which is also discussed in **Supplementary Material S4**. Here in the main text, we report quantifications for ROIs defined in our own ‘*splitting’* scheme, wherein we delineate between 7 core and belt regions based on the presence of tonotopic reversals ^22,30^, variations in tuning width ^28,31^ (here defined as FWHM), degree of speech selectivity ^27,32^ and myelination ^33^. For completeness, in **Supplementary Material S5 and Supplementary Figures S6-S9**, we provide equivalent quantifications for ROIs defined in two alternative schemes: i) a simple *‘lumping’* scheme that binarizes the tonotopic population of vertices into ‘core’ and ‘non-core’ regions and ii) a *‘neutral’* scheme, that ignores our own model parameters and uses the existing HCP multi-modal parcellation of auditory cortices ^61^.

### Bootstrapping Procedure

*p*-values were computed using 10^4^ fold bootstrap procedures, with the estimation of parameters (mean differences, regression slopes) for each bootstrap sample weighted by out-of-sample R^2^. To test whether bootstrapped distributions differed from a certain threshold, *p*-values were defined as the ratio of bootstrap samples below versus above that threshold multiplied by 2 (all reported *p*-values are two-tailed). All *p* values were then corrected for multiple comparisons across ROIs using a false discovery rate correction. For any comparisons wherein all bootstrapped samples fell on one side of the threshold, the *p* value was coded as 1/10^4^. For presentation, we -log^10^ transformed *p* values resulting from a positive difference between a given reference and comparison ROI (resulting in positive values) and log10 transformed *p* values resulting from negative differences (resulting in negative values). The same transformation was applied to *p* values resulting from positive v negative associations between variables. All reported *p* values thus reside on a scale between -4 and 4.

## Acknowledgements

Data were provided by the Human Connectome Project, Washington University - Minnesota Consortium (Principal Investigators D. Van Essen and K. Uǧurbil; 1U54MH091657) funded by the 16 NIH Institutes and Centers that support the NIH Blueprint for Neuroscience Research, and by the McDonnell Center for Systems Neuroscience at Washington University in St. Louis. NH was supported by a Leverhulme Trust Early Career Fellowship (ECF 2019-305) that funded his time on this project.

## Data Availability

Source data for this article are freely available after registration to the HCP website (https://www.humanconnectome.org/). We have provided our estimated model parameters and ROI definitions in ‘gifti’ surface format (both in original HCP 59k space and upsampled to ‘fsaverage’ 164k space) in the following figshare repository: https://figshare.com/projects/Data_from_Naturalistic_Stimulation_Reveals_the_Topographic_Organization_of_Human_Auditory_Cortex_/117288. We also provide the HCP ‘pycortex’ subject that was used to produce all visualisations.

## Code Availability

Code for performing model fitting is available at: https://github.com/N-HEDGER/Tonotopy_2021

