## Supplementary Material for "Naturalistic Audiovisual Stimulation Reveals the Tonotopic Organization of Human Auditory Cortex"

**Supplementary Material for ‘Naturalistic Audiovisual Stimulation Reveals the Topographic Organization of Human Auditory Cortex’**

**S1. Predictive Capacity of the CSS model.**

**S2. Stability of Parameter Estimates.**

**S3. PCA Analysis of ‘Early’ and ‘Late’ Subjects.**

**S4. Strategies to Delineate ROIs.**

**S5. Quantifications for ‘Neutral’ and ‘Lumping’ Schemes.**

**S6. Null Model and Vertex Selection**

**S7. Spectrograms of Analysed Movie Sequences.**

**S8. Alternative Quantification of pRF Size.**

### Supplementary Material S1. Predictive Capability of the CSS Model

#### Comparison to Fixed Exponent Model

We compared the cross-validated performance of our CSS model (which estimates n as a free parameter) to a constrained model with a fixed exponent of .3 (the estimated mean of *n* values). These data are depicted in **Figure S1F** and **S1G**. The CSS model outperformed the constrained model in 94% of vertices. Interestingly, the spatial profile of model improvement indicates the smallest benefits within core regions along HG (dark region of **Figure S1G**) - pointing to the relative importance of this parameter in explaining responses in non-primary regions.

#### rRMSE

As a further assessment of the predictive capacity of the model, we calculated the rRMSE metric as follows:

$$rRMSE = \frac{(RMSE(M1, D2) + RMSE(M2, D1))}{2(RMSE(D1, D2))}$$

Where D1 and D2 are split halves of the data, M1 and M2 are the model predictions for the corresponding data. This metric thus references the generalization performance of the model against test-retest reliability of the underlying data. Accordingly, values of >1 indicate that model predictions are less accurate than test-retest reliability. This data is depicted in **Figure S1H** and **S1I**. The median rRMSE value was 1.69, with substantial variation throughout the tonotopic population (range= 0.83-3.13). One interesting aspect of the data revealed by **Figure S1H** is that the model was increasingly limited in its ability to capture data towards STG, supporting the notion that the responses in these regions are less yoked to basic spectral tuning.

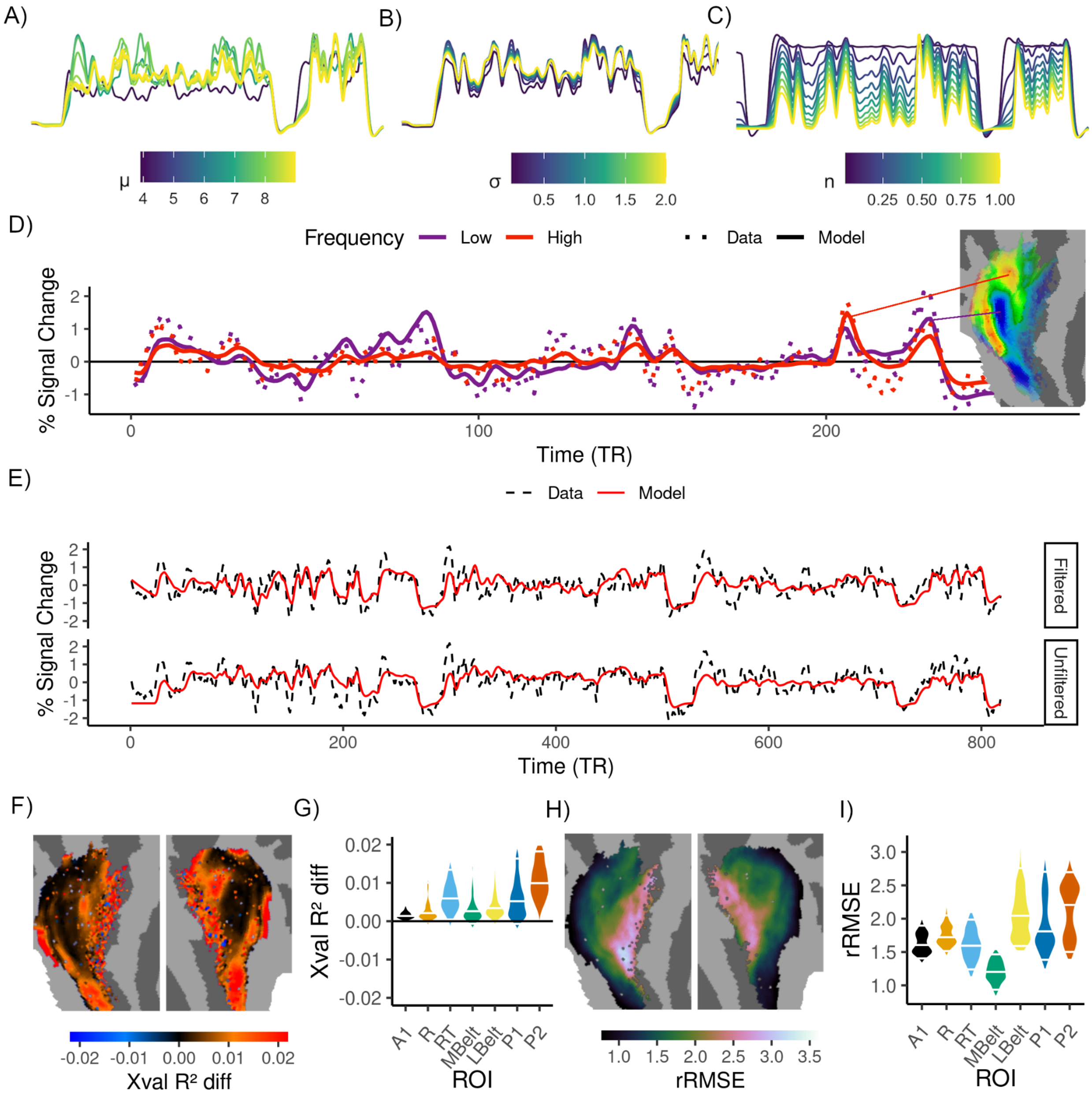


***Figure S1.*** Panels **A-C** depict a set of model predictions derived from varying the μ, σ and n parameters of the CSS model, as depicted by the colorbars*.* To visualise the effect of uniquely varying each parameter, the predictions in each plot are generated by fixing the other two parameters. All predictions are peak-normalised. **D)** Depicts data and model fits for representative vertices that prefer low and high auditory frequencies whose locations are indicated on the flatmap. Panel **E** depicts representative data and model predictions that are filtered and unfiltered using the Savitzky Golay filter described in the method section. The data corresponds to the first movie. **F-G)** Depicts the difference in cross-validated model performance between the CSS model and a constrained model with a fixed exponent Positive values/ red regions indicate superior performance of the CSS model. **H-I)** Depict the rRMSE throughout the tonotopic population. Violins are normalised to have equal maximum width. The 5th, 50th and 95th percentiles of the distributions are demarcated by white vertical lines.

**Supplementary Material S2. Stability of Parameter Estimates.**

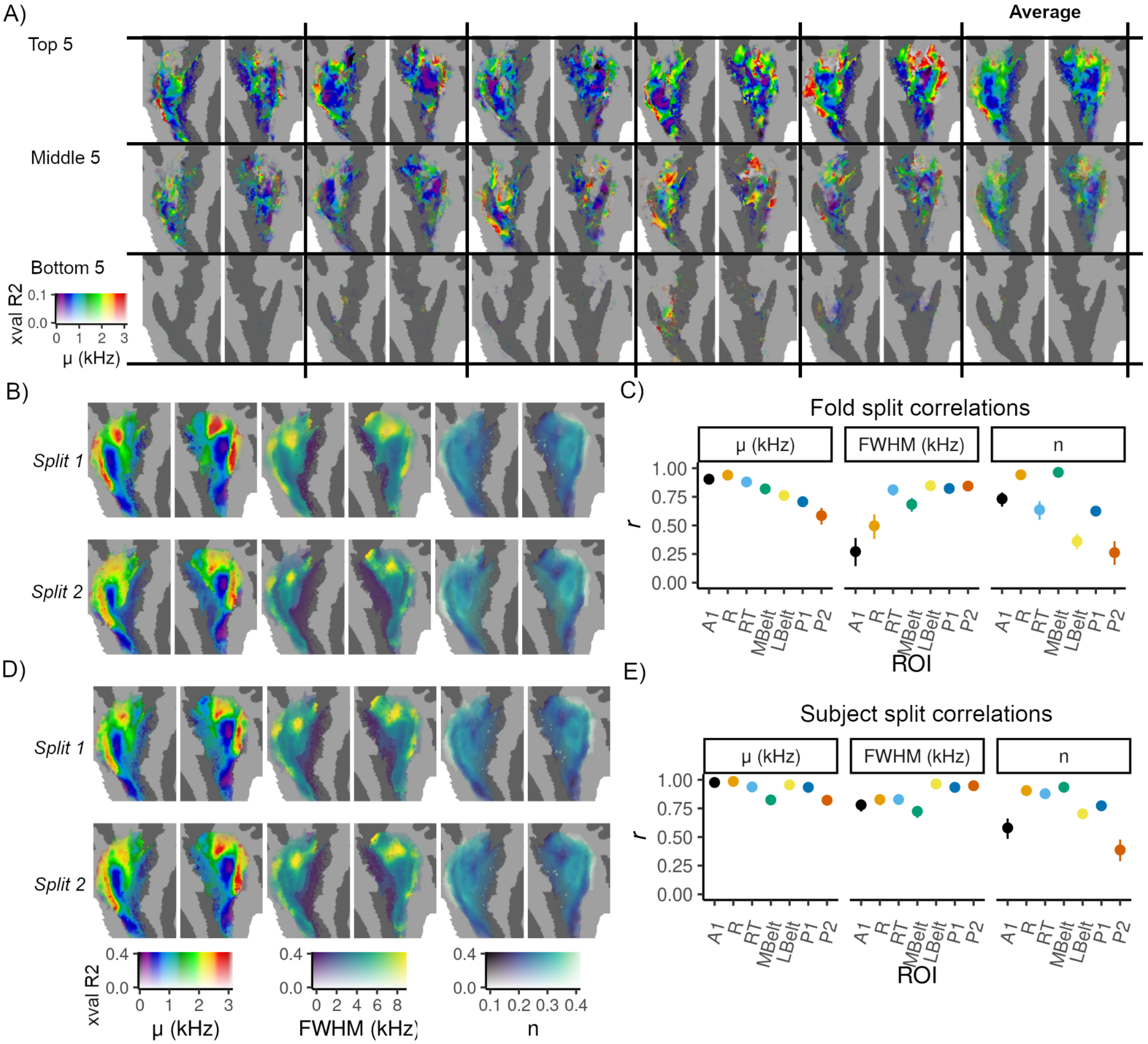


***Figure S2.*** **A)** Depicts preferred frequency (μ) data for the top, middle and bottom 5 subjects and the associated averaged estimates (see **Methods** for participant ranking). The 2D colormap depicts μ across the x axis and generalisation performance (*Xval* $R^2$) defined by transparency across the y axis. The data from the top 5 subjects exhibit features of the tonotopic arrangement revealed from the aggregated data. (Figure 2B in the main text). These patterns begin to break down and are less discernible in the middle 5 subjects. Essentially no tonotopic arrangement is visible in the bottom 5 subjects, reflecting the poor cross-validation performance. These results outline that tonotopic maps are not easily identified based on individual subject data. **B)** Depicts flatmaps for the μ, FWHM and n parameters across two fold splits of the data. **C**) Depicts the corresponding correlation coefficients between these splits as a function of ROI. Error bars indicate the 95% confidence interval across vertices. **D-E)** Depicts the same as **B-C** across two independent subject split-halves of the data. Across-split associations were detected between all parameters in all ROIs (all $p < .001$, all $r > .26$), indicating the stability of parameter estimates across and within participants.

**Supplementary Material S3. PCA Analysis of 'Early' and 'Late' Subjects.**

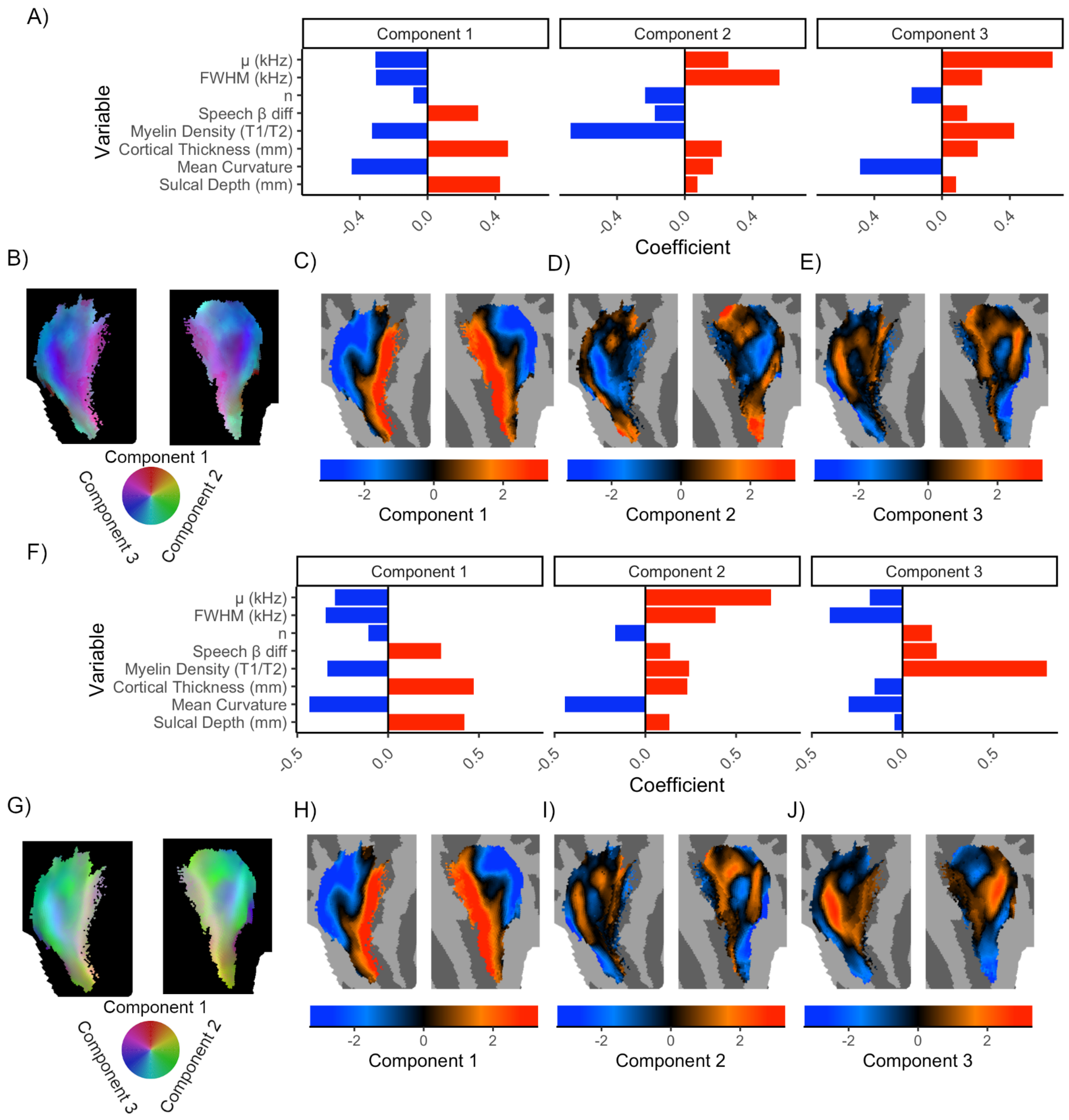


***Figure S3.*** Panels **A-E** depict the PCA results for the 'early' subject **A)** Depicts the feature loadings of each parameter onto 3 principle components. **B)** Shows the components in RGB color space. Panels **C-E** depict the individual components. Panels **F-J** depict the same data for the 'late' subject. There is a high level of correspondence between the PCA components revealed for the two participants. Note that component 2 for the early subject (panel **D**) is essentially the inversely-signed equivalent of component 3 for the late subject (Panel **J**) and that component 3 for the early subject (panel **E**) is essentially component 2 of the late subject (Panel **I**) .

### Supplementary Material S4. Strategies to Delineate ROIs.

#### Qualifications Regarding ROI Definitions.

As a proviso, please note that we do not wish to claim that the ROIs we define here reflect 'cortical field maps' - a status for which they would need to be defined by two orthogonal dimensions of a fundamental sensory dimension (e.g. tonotopy and periodicity) [1,2]. Since our ROIs are primarily defined by tonotopy, they could be thought of as a set of 'cortical gradient maps' [1] that we define here to provide some spatial precision to our quantifications. Although we note that the ROIs we define have a good correspondence to the 'working model' of auditory cortex proposed by [3] and also to the HCP parcellation of auditory cortex [4], we are agnostic about whether they constitute distinct regions responsible for distinct sensory computations. Secondly, we do not wish to imply any direct correspondence between the ROIs we define here and those in the Macaque models of auditory cortex [5], particularly those defined beyond core regions. To reflect this, we use altered terminology (e.g. '*MBelt*' instead of '*Medial Belt*') similar to that used by [4].

#### 'Splitting' Scheme

ROIs were drawn using pycortex [6]. In drawing these ROIs, we adhered to the following guiding principles: i) Each region must be sensitive to structured auditory information (i.e. it must be part of the sub-population of tonotopic vertices we identify from the procedure outlined in **Supplementary Material S6**. ii) Each region must contain a representative, spatially non-repeating range of the preferred frequencies in the sub-population. iii) Each region must be identifiable in both hemispheres of both the 'early' and 'late' subjects. iv) Finally, the drawing of ROIs must be guided by using the 'working model' of human auditory cortex as outlined by [3] as a 'prior' for estimating boundaries between regions. As indicated in the main text (see **Data Access Statement**), we provide a pycortex subject database entry and our estimated model parameters for the HCP data, facilitating further, alternative ROI schemes to be drawn.

A clear set of tonotopic reversals could be discerned along HG (**Figure S4A**). Within the medial two-thirds of HG, there is a tonotopic gradient that proceeds from high frequencies (**i**) in a roughly antero-lateral direction towards a low frequency minimum where a tonotopic reversal can be observed (**ii**) Further anterior, there is a reversal at a high frequency maximum (**iii**). Further anterior still, there is an additional reversal at low frequencies (**iv**), beyond which no clear tonotopic progression can be observed. Taken together, this resembles the high-low-high-low tonotopic pattern that is characteristic of the core fields *A1, R and RT* [7]. Guided by the location of these frequency reversals, we defined initial boundaries between core fields *A1, R and RT* (**Figure S4A**). To delineate the medial and lateral boundaries of this 'core' region, we first drew a boundary more medially at a region marked by an abrupt increase in FWHM (**Figure S4B**), consistent with the observation that the auditory core is flanked by regions with higher bandwidths [8]. More laterally, FWHM did not appear to serve as an informative cue for drawing a boundary. Instead, we drew the lateral boundary at a region that was marked by an abrupt increase in speech selectivity (**Figure S4C**), consistent with the observation that speech selectivity is reliably a property of non-core auditory cortex [9,10]. Interpolating between these two boundaries reveals that our definition of this 'core' agrees very well with the region of highest myelin density (**Figure S4D**), which has been a previously reported property of the auditory core, and of primary sensory cortices more generally [11,12].

Laterally to the core, additional frequency reversals could be discerned (**Figure S4E**). We drew an additional ROI extending laterally from the core to the high frequency maximum at **v**). This region, which contains its own, single gradient that mirrors that of *A1*, we labelled as '*LBelt*'. Notably, the postero-medial section of the *LBelt* that adjoins *A1* is marked by a tonotopic reversal at low frequencies and contains the majority of the tonotopic gradient (**Figure S4F**). Moreover, the more anterior section that neighbours *R* and *RT* is heavily biased towards lower frequencies and is characterised by high speech selectivity (**Figure S4G**). These properties are all in good agreement with the properties of the lateral belt outlined by Moerel and colleagues [3].

Extending postero-laterally to LBelt, we drew an additional ROI that terminated at a further, more subtle tonotopic reversal at low-frequencies (**vi**). We labelled this region as '*P1*'. From the posterior boundary of this region, we drew one final ROI that appeared to contain an additional, but less salient, tonotopic gradient up until where the tonotopic population terminated (**Figure S4H**). We labelled this region '*P2*'. We use unique terminology (*P1* and *P2*) to refer to these regions because regions beyond the lateral belt (i.e. the parabelt) are not traditionally defined by tonotopic reversals. A reversal at low frequencies could be discerned around STG (**vii**) which served to define a dorsal boundary for all ROIs placed laterally to the core. All tonotopic regions positioned medially to the core were defined as '*MBelt*'. The resulting ROIs are depicted in **Figure S4I**.

**'Lumping' Scheme**

For the lumping scheme, we used the definition of 'core' regions defined in the splitting scheme (*A1, R, RT*) and collapsed them into a single 'core' region. The remaining regions (*MBelt, LBelt, P1, P2*) were defined as 'non-core' (**Figure S4J**).

**'Neutral' Scheme**

For this final scheme, we used an existing multi-modal parcellation of cerebral cortex [4]. From this, we drew all 5 regions defined as 'primary auditory' *(A1, LBelt, MBelt, PBelt RI*) and all 8 regions defined as 'auditory association' cortices (*A4, A5, STSdp, STSda, STSvp, STSva, STGa and TA2*). This parcellation is depicted in **Figure S4K**. The definition of these regions were primarily based on factors such as connectivity with the medial geniculate nucleus, myelination, cortical thickness and speech selectivity. A detailed description of these regions is provided in the supplementary material of [4] .

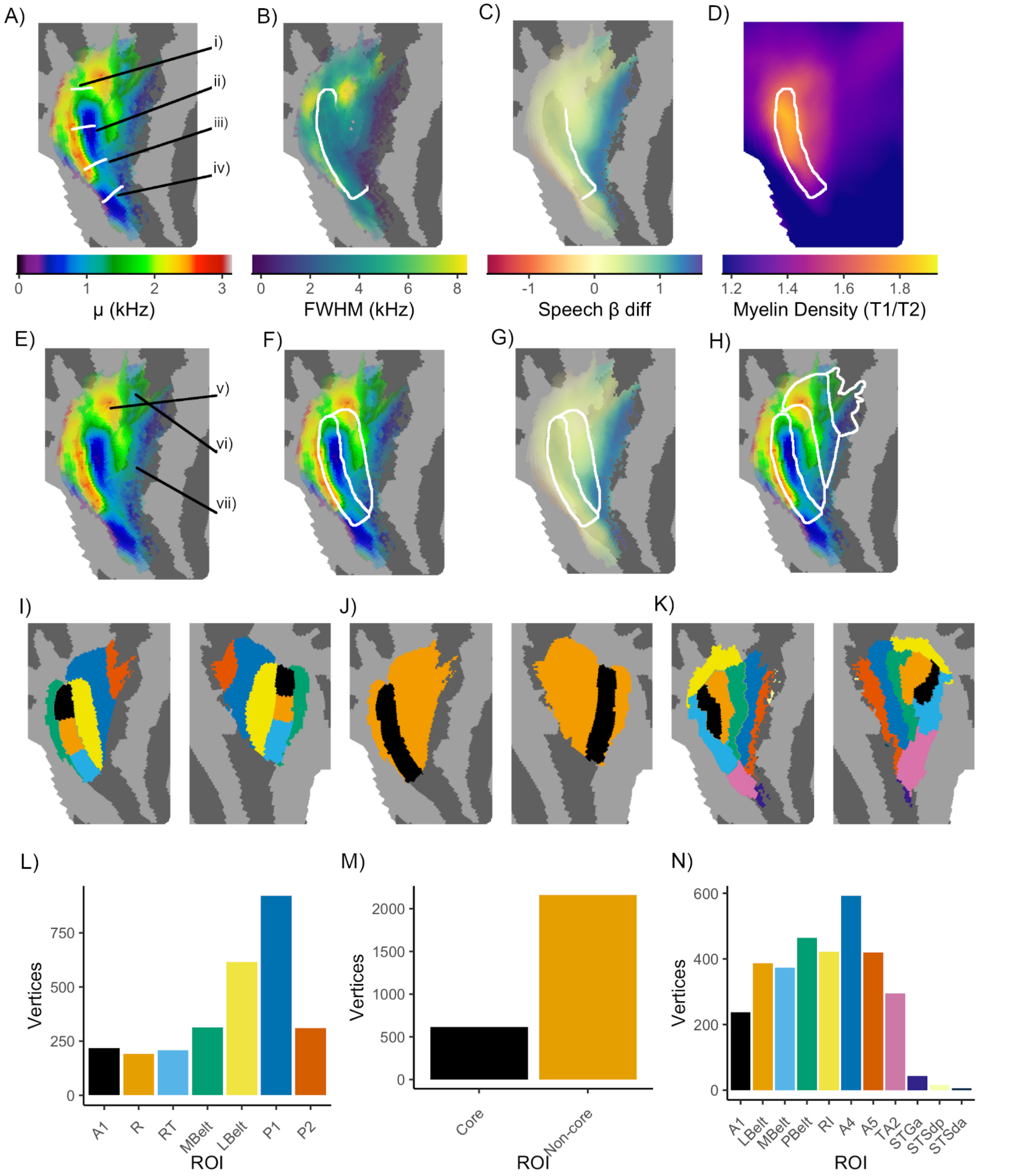


***Figure S4.*** Panels **A-H** depict information sources that were used to define different ROIs. Preferred frequency (**A**). FWHM (**B**) Speech-selectivity (**C**). Myelin density (**D**). Preferred frequency (**E-F**). Speech-selectivity data (**G**). Preferred frequency (**H**). White lines are used to indicate boundaries that were guided by these information sources, as described in the above text. Panels **I-K** show the definition of ROIs for the splitting, lumping and neutral schemes respectively. Note that the data are masked to depict only the vertices in the tonotopic population we analyze in the main text. Different colors delineate different regions, whose names and the associated number of vertices are given in panels **L-N**.

**Location of Analysed Vertices.**

To reference the subset of tonotopic vertices we analyse in the main text (see **Supplementary Material S5**) against an existing benchmark, it is first informative to relate their position to the ROIs defined in the neutral HCP parcellation. **Figure S5C** and **S5F** illustrate that all ROIs defined as ‘primary auditory cortex’ in the neutral HCP parcellation (i.e. *A1, LBelt, Mbelt, PBelt, RI*) reside within our tonotopic population. Interestingly, there is also near-complete overlap with several ROIs defined as ‘auditory association cortex’ (*A4, A5, TA2*) - although some of the more ventral ‘association’ regions are barely represented (*STGa, STSda, STSdp*) and the ventral portions of the STS (*STSva, STSvp*) are not represented at all.

**Relations Between the ‘Splitting’ and ‘Neutral’ ROI scheme.**

**Figure S5G** depicts the overlap between ROIs defined in our splitting scheme and those in the neutral HCP scheme. Overall there is a good level of consistency between the two, but there are potentially informative differences to observe. First, although there is generally good agreement between our definitions of *A1*, the *neutral* definition also overlaps with our definition of the core field *R*. Since the *neutral* parcellation did not incorporate tonotopy data, the *R* field (whose location is usually determined by tonotopic reversal along HG) was not defined. It therefore makes good sense that the *neutral* definition of *A1* overlaps with our definition of *R*. Inspection of **Figures S5A** and **C** reveal that the *neutral* regions lateral to *A1* (*LBelt, PBelt, A4 and A5*) are aligned more with STG, whereas the regions that we define laterally to the core (*LBelt, P1, P2*) are larger and aligned more with HG. Functionally, the HCP parcellation was partially based on data from language-sensitive task contrasts (e.g. the difference in activity to listening to a story vs performing math problems), but not tonotopy. As such, *neutral* ROIs were most likely biased primarily along the axis of increasing speech selectivity (aligned with STG), whereas our ROIs are more aligned with the axis of the tonotopic gradients observed in the data (aligned with HG). In practice, this entails that our ROIs lateral to the core span multiple *neutral* regions (**Figure S5G**). Finally, the *neutral* ROI defined as *STGa* is not represented in our scheme, as we failed to detect any clear tonotopic progressions anterior to what we define as *RT*.

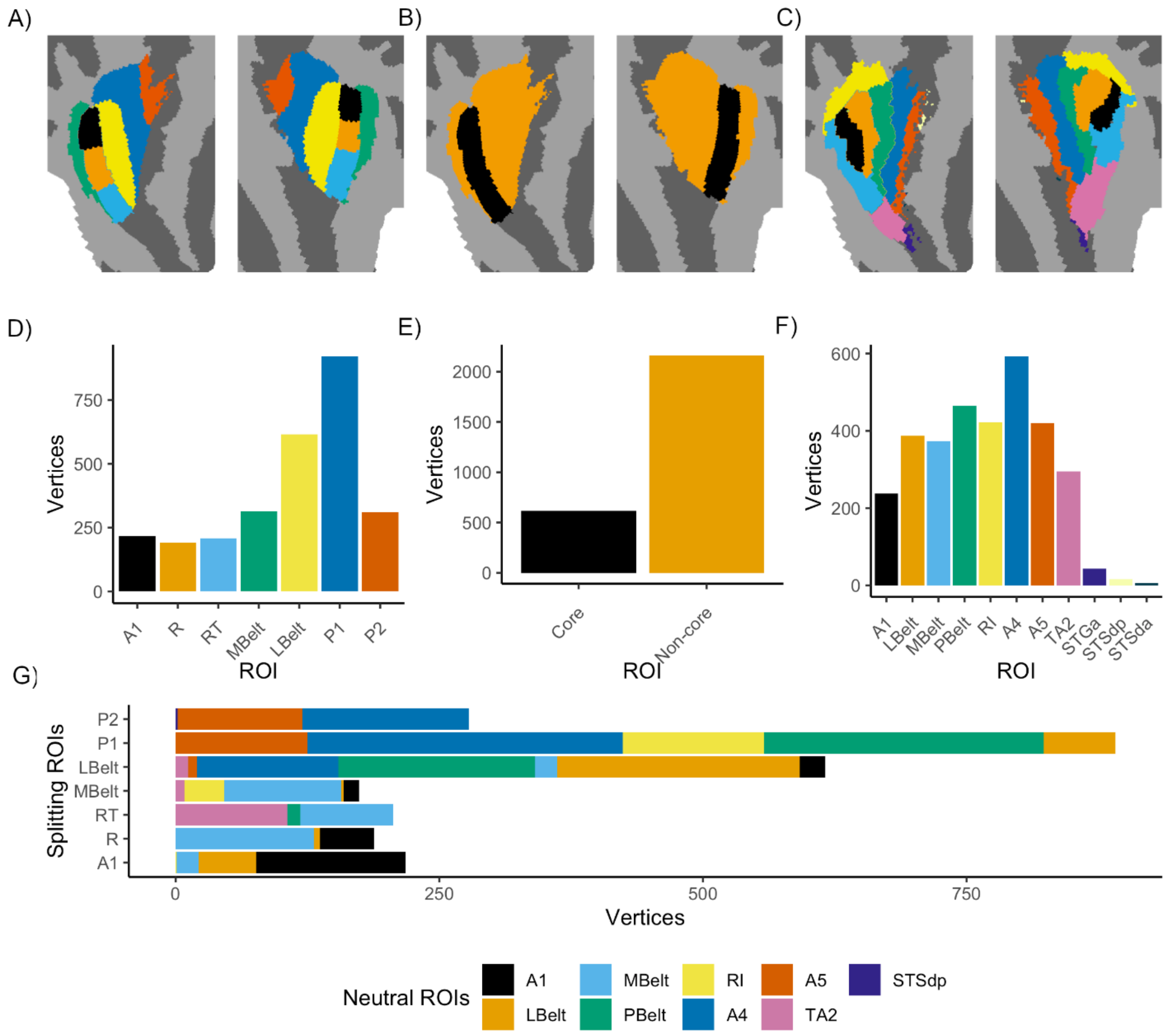


***Figure S5.*** Panels **A-C** show the definition of ROIs for the splitting, lumping and neutral schemes respectively. Note that the data are masked to depict only the subset of tonotopic vertices we analyse in the main text. Different colors delineate different regions, whose names and the associated number of vertices are given in panels **D-F**. Panel **G** depicts the overlap between regions defined in the splitting and neutral schemes.

**Orientation of the Core Fields.**

All of the ROI definitions here (including the HCP parcellation) include 'core' regions that run along HG. We must therefore acknowledge that there is also controversy regarding the orientation of the auditory 'core' with respect to HG. Some sections of the auditory neuroscience community prefer an arrangement that runs across HG ('orthogonal' interpretation), rather than along HG ('classical' interpretation) [7]. This dispute results to a large extent from the fact that the single dimension of tonotopy alone may be deficient for resolving these two competing possibilities [1,3]. We note however, that the more 'classical' scheme we define here was also guided by multi-modal data (variations in myelin, FWHM and speech selectivity). Moreover, this classical organisation also agrees well with a recently refined cytoarchitectonic atlas of auditory cortex that defines 3 regions (Te 1.1, Te 1.0 and Te 1.2) running in a 'classical' orientation along HG [13]. In turn, these 3 regions overlap with independent myeloarchitectonic quantifications that revealed 'keyhole' shaped region of high myelin density elongated posteromedially to anterolaterally along HG [14] - similar to that depicted in **Figure 3E** and **3M** in the main text.

**Supplementary Material S5. Quantifications for 'Neutral' and 'Lumping' Schemes.**

**Neutral Scheme**

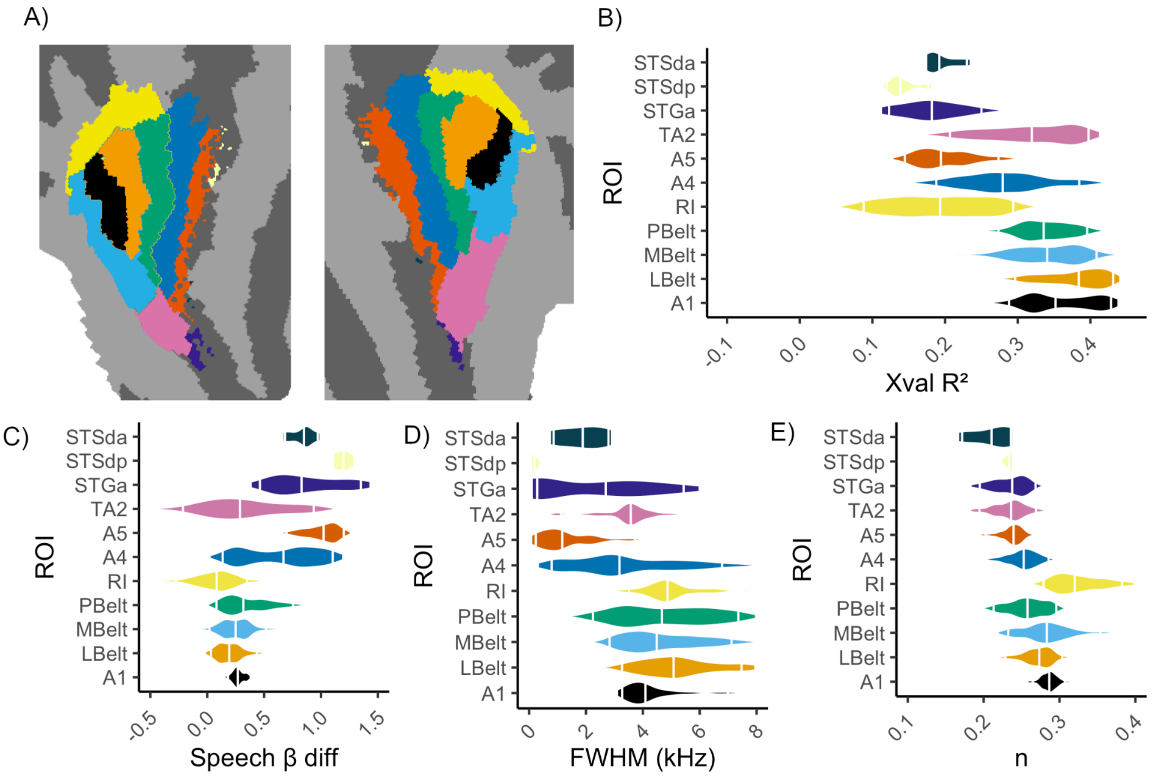


***Figure S6.*** **A)** Key to show the definition of ROIs for the 'neutral' scheme. These correspond to the colours of the violins in the remaining figures. Panels **B-E** show the per-ROI distributions for variance explained by the CSS model (**B**), speech selectivity (**C**), FWHM (**D**) and n (**E**).

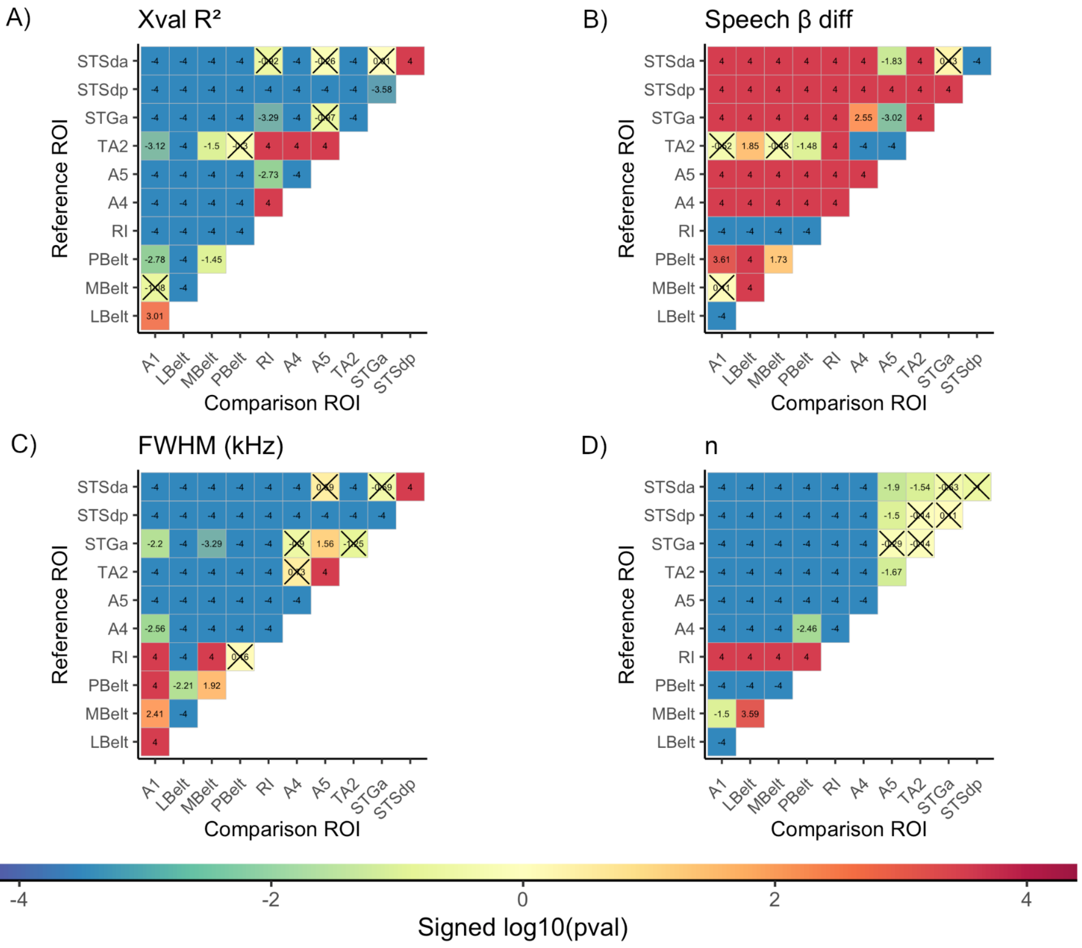


***Figure S7.*** Depicts the between-ROI pairwise differences for the variance explained by the CSS model (**A**), speech selectivity (**B**), FWHM (**C**) and n (**D**). The values in each cell are signed log10 transformed *p* values. Larger values for a parameter in a reference ROI (y axis) compared to a comparison ROI (x axis) are assigned positive values (p values are -log10 transformed). Smaller values in the reference ROI are assigned negative values (*p* values are log10 transformed). *p* values that do not reach the false discovery rate corrected alpha level of .05 are marked by a cross in the corresponding cell. The corresponding colorbar for all *p* values can be found at the bottom of the figure. We advise due caution in interpreting *p* values involving ROIs within which very few vertices were analyzed (e.g. *STGa, STSdp, STSda* - **Figure S4F**) due to the limited number of unique bootstrapped samples that can be drawn.

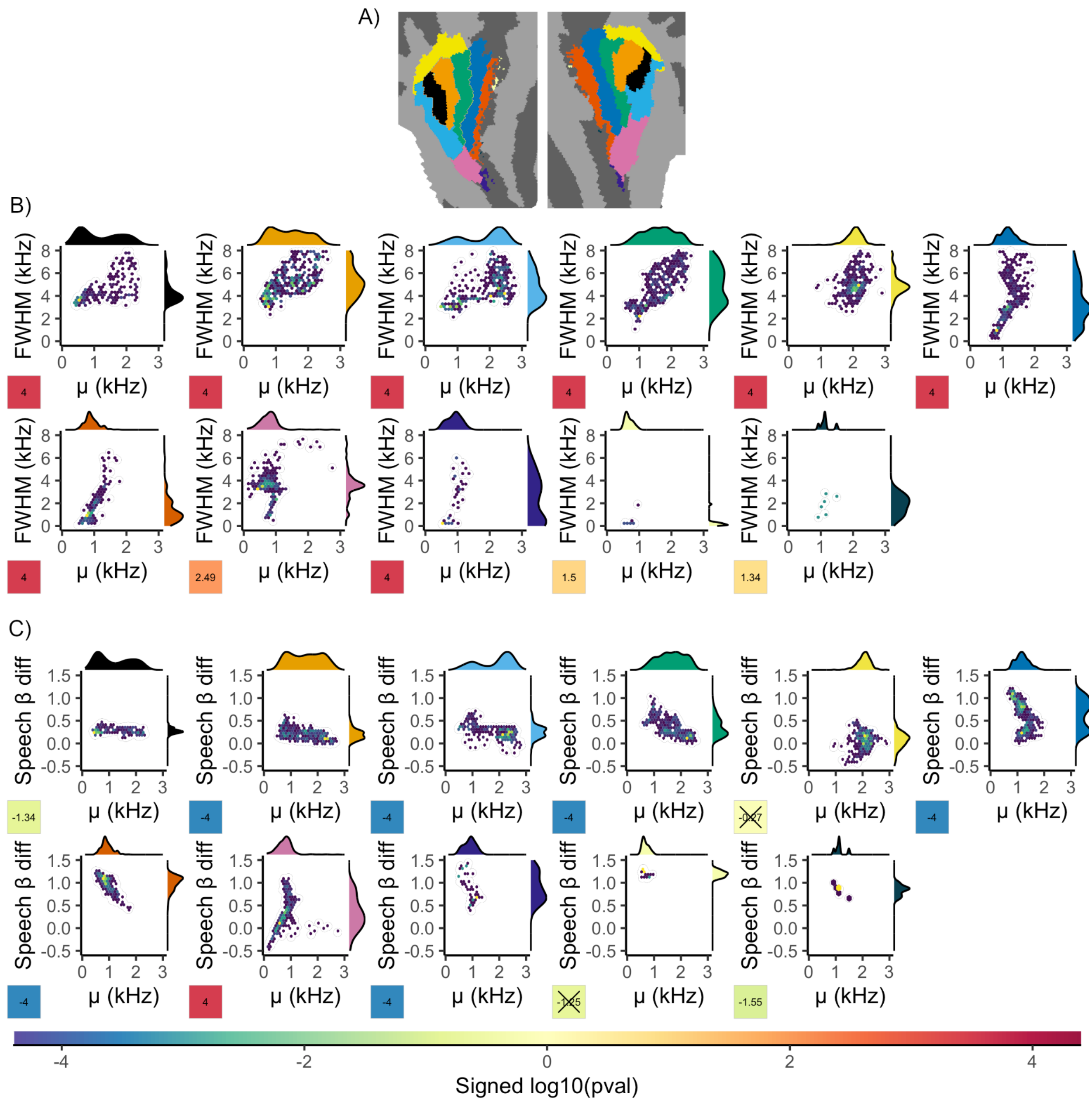


***Figure S8.*** **A**) Key to show the definition of ROIs for the 'neutral' scheme. ROI color corresponds to the colour of the marginal density plots in the remaining figures. Panel **B** shows hexbin plots for the relationship between μ and FWHM in each ROI. The cells in the bottom left of each plot depict the signed log10 transformed *p* values (positive correlations have positive values). Panel **C** shows hexbin plots for the relationship between μ and speech selectivity.

**Lumping Scheme**

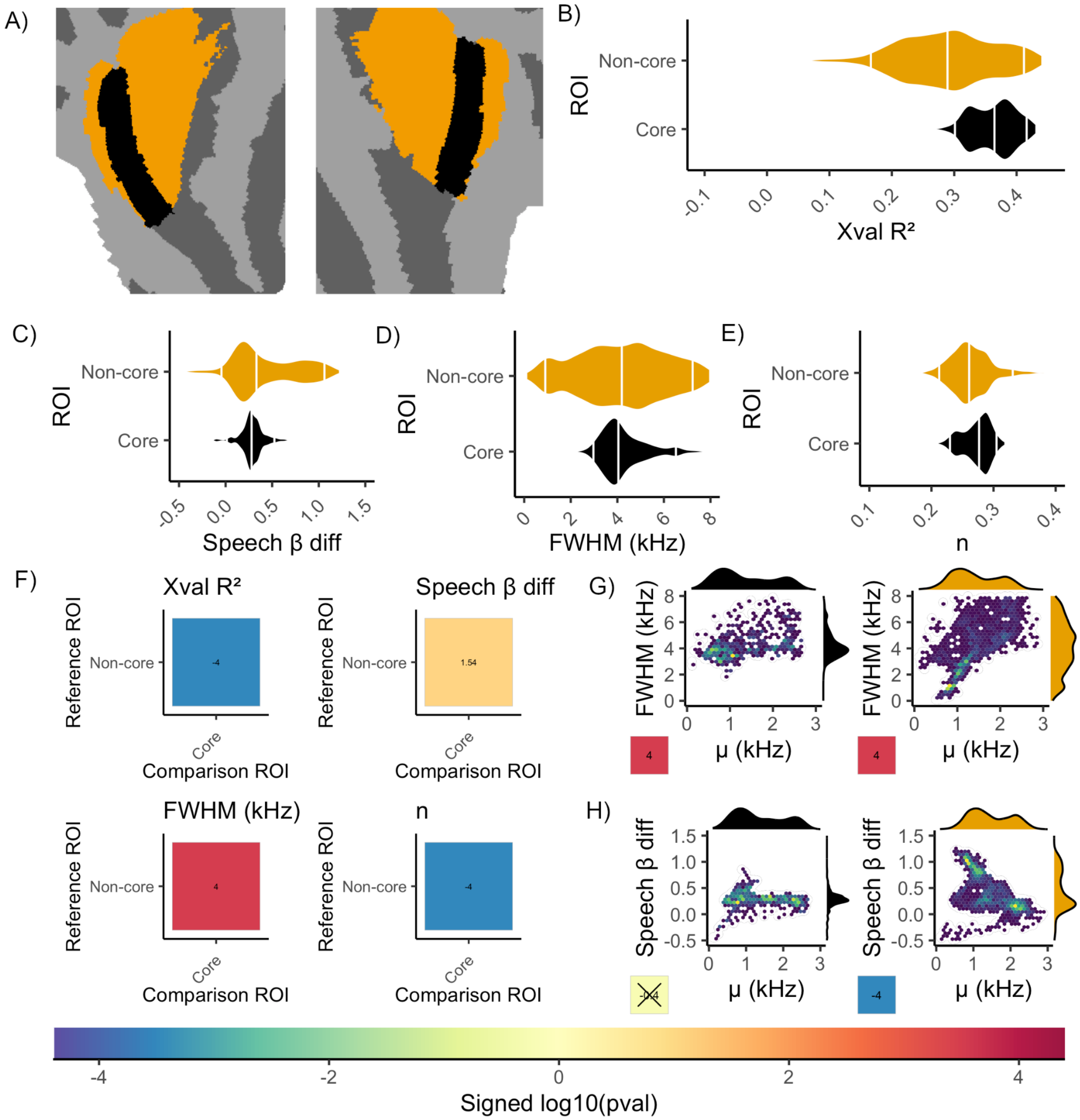


***Figure S9.*** **A)** Key to show the definition of ROIs for the 'lumping' scheme. ROI color corresponds to the colours of the violin /marginal density plots in the remaining figures. Panels **B-E** show the per-ROI distributions for variance explained by the CSS model (**B**), speech selectivity (**C**), FWHM (**D**) and n (**E**). Panel F depicts the between-ROI pairwise differences for the corresponding variables depicted in panels **B-E**. Panel **G** shows hexbin plots for the relationship between μ and FWHM in each ROI. Panel **H** shows hexbin plots for the relationship between μ and speech selectivity.

### Supplementary Material S6. Null Model and Vertex Selection.

**Figure S10A** depicts the out of sample variance explained (xval $R^2$) by the CSS model across the entire cortical surface. Consistent with prior studies, the best performance is observed in a large expanse of the superior temporal cortex. Notably, there is also good performance visible in a large expanse of occipital and fronto-parietal cortices. **Figure S10B** depicts the estimated exponent (n) parameter. Notably, in temporal cortex, the estimated exponent values are close to 0.3, which is in good agreement with the compressive power function performed on auditory signals by the cochlea [15]. Moreover, this parameter also clearly differentiates temporal cortex from the occipital cortex, where highly-compressive, near-zero exponent values are observed. We therefore suspected that such pRF estimates remote from temporal cortex capture non-tonotopic responses that reflect total stimulus energy or generic 'task-positive' responses that would be well approximated by a CSS model with a very low exponent.

Subsequently, a non-tonotopic 'sound-on' null model was used to identify and exclude vertices that responded to any sound that was presented. Specifically, the out-of-sample variance explained by this null model was subtracted from the CSS model. Confirming our suspicion, this had a robust, deleterious impact on occipital and frontal cortices (Figures **S10C- S10E**). Critically, however, it clearly reveals a spatially contiguous population of vertices sensitive to structured auditory information in superior temporal cortex (**Figure S10E**, white outline). Only vertices for which the CSS model variance explained exceeded the null model by more than 5% were entered into subsequent analyses.

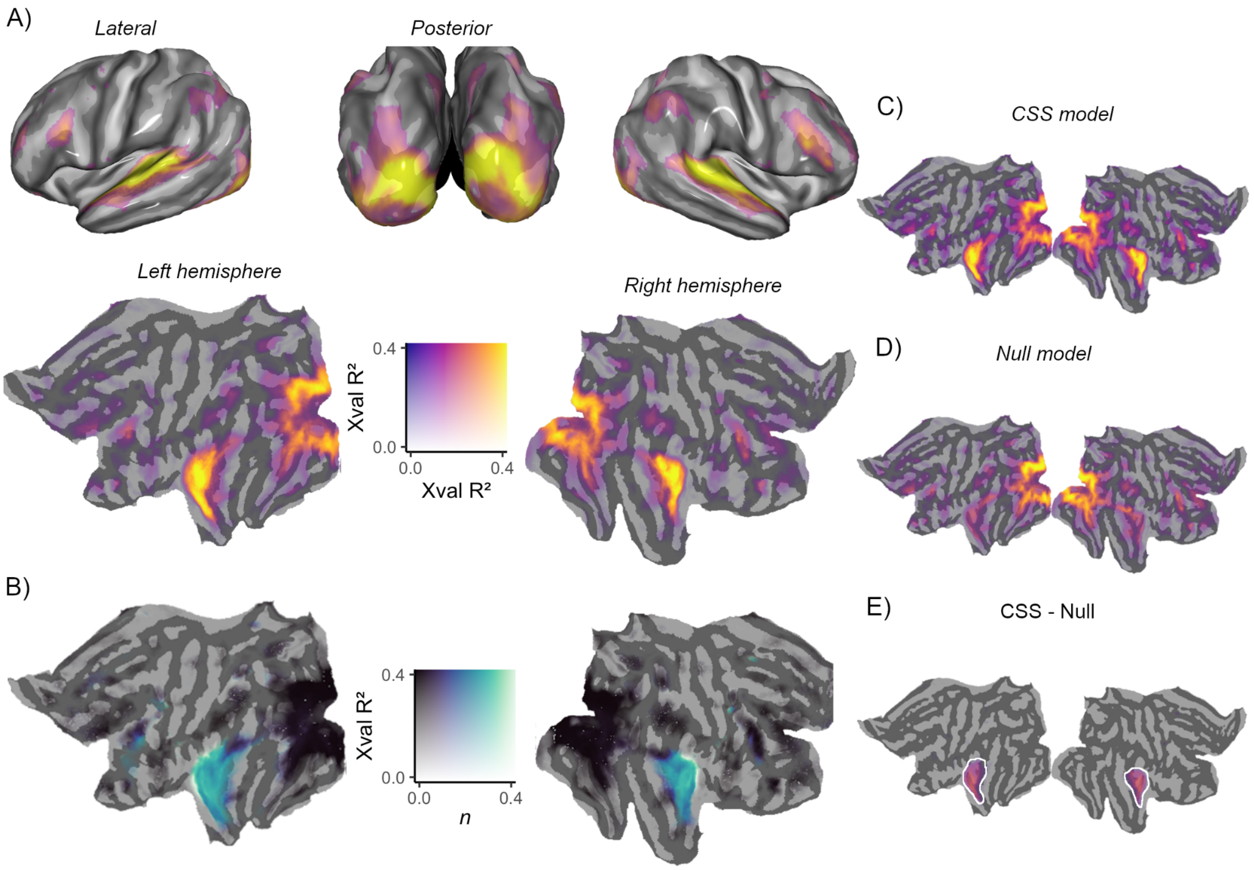


***Figure S10.*** **A**) Depicts cross validated variance explained by the CSS model. **B**) Depicts the exponent parameter *n*. The auditory cortex, (turquoise) is characterised by high n values and is clearly distinguishable from the visual cortex, which is characterised by near-zero n (black). Panels **C-E** depict the CSS model performance (**C**), null model performance (**D**) and the tonotopic population revealed by subtracting null model performance from CSS model performance (**E,** white outline).

**Supplementary Material S7. Spectrograms of Analysed Movie Sequences.**

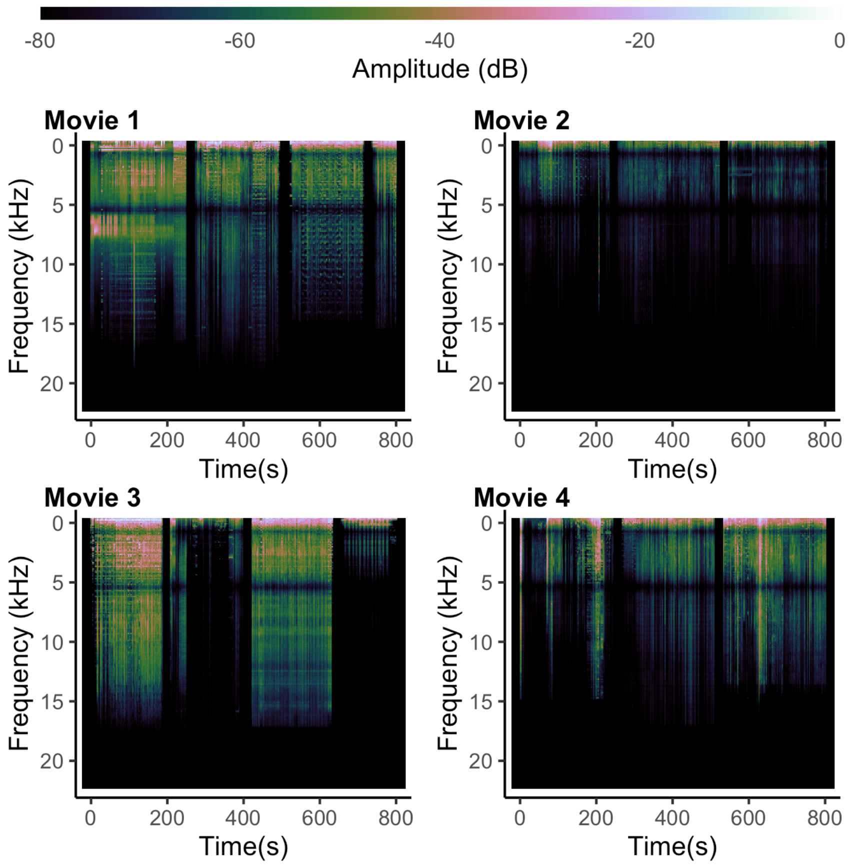


***Figure S11.*** Displays spectrogram representations of the 4 movie sequences analysed. Mean sound intensity (expressed in decibels (dB)) is presented as a function of frequency in KHz and time in seconds (corresponding to a TR). Amplitude values are normalised such that the reference value of 0 demarcates the peak amplitude across all movie sequences.

**Supplementary Material S8. Alternative Quantification of pRF Size**

**Figure S12** depicts estimates of pRF size expressed in octaves. These were defined by calculating the lower (*f1*) and upper (*f2*) frequencies at which the pRF function reached half of its maximum and subsequently converting via:

$$log2(f2/f1)$$

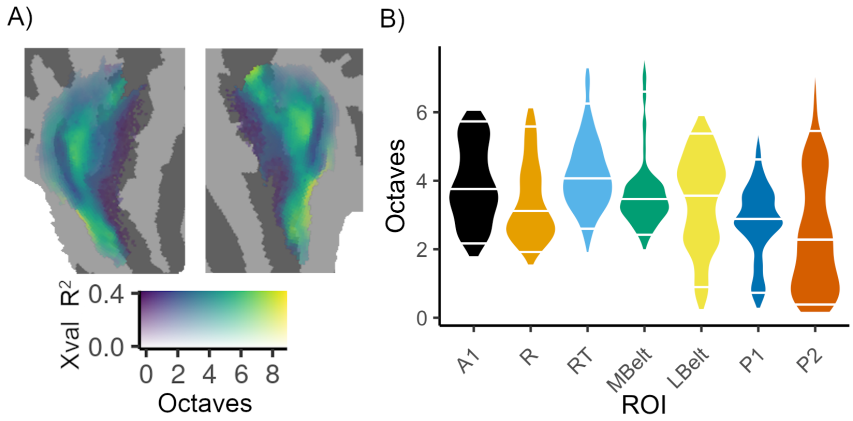


***Figure S12.*** **A)** Flatmap depicting pRF size, expressed in octave units. **B)** Per-ROI distributions of octave estimates.
